# Regulation of the Yolk Microtubule and Actin Cytoskeleton by Dachsous Cadherins during Zebrafish Epiboly

**DOI:** 10.1101/2025.05.10.653271

**Authors:** Gina Castelvecchi, Linwei Li, Jimann Shin, Nanbing Li-Villarreal, Isa Roszko, Paul Gontarz, Tiandao Li, Bo Zhang, Diane Sepich, Lila Solnica-Krezel

## Abstract

Epiboly is a crucial morphogenetic process during early animal embryogenesis that expands surface area of embryonic tissues while thinning them. During zebrafish development, epiboly spreads the superficial enveloping layer (EVL), germ layers, and yolk syncytial layer to cover the yolk cell. Here we investigated functions of the three zebrafish *dchs* genes, *dchs1a, dchs1b and dchs2* that encode large atypical cadherins and report that they have partially overlapping functions in epiboly progression. We have inserted GFP at the C-terminal Dchs1b intracellular domain of the endogenous *dchs1b* locus using homologous recombination. We observed the resulting Dchs1b-GFP fusion protein localized in both the cell membrane and the cytoplasm of EVL and embryonic cells during gastrulation. The dynamic microtubule and actin cytoskeleton of the yolk cell are essential for epiboly. Our studies of the yolk microtubule network demonstrate that these microtubules are more bundled and show faster polymerization during epiboly in *dchs* triple loss-of-function mutant embryos than in wild-type embryos, indicating that *dchs* genes are required for limiting microtubule polymerization and promoting dynamics during epiboly. The epiboly progression defects of *dchs1b* deficient mutants were suppressed by mutations in the *tetratricopeptide repeat protein 28* (*ttc28)* gene encoding a cytoplasmic protein previously shown to bind to Dchs1b intracellular domain and alter microtubule dynamics during early cleavages. We further demonstrate that MZ*dchs1b* mutants exhibit abnormal organization and dynamics of yolk cell actin cytoskeleton during epiboly. Together, these lines of evidence as well as our transcriptomic analyses support the notion that like during early embryonic cleavages, Dchs1b plays a major role, while Dchs1a and Dchs2 proteins have supporting roles in regulating microtubule dynamics and organization of both microtubule and actin cytoskeleton to ensure normal epiboly.

## Introduction

Epiboly is one of the major, evolutionarily conserved morphogenetic processes during animal development that involves expansion and concurrent thinning of the embryonic tissues. In zebrafish, epiboly is well studied and initiates just after zygotic genome activation (ZGA) (Bruce and Heisenberg, 2020). At this stage, the zebrafish embryo is comprised of three cellular compartments. The epithelial enveloping layer (EVL) consisting of the most superficial cells is attached at its edge to the yolk cell. Sandwiched between them are mesenchymal deep cells that will give rise to the embryo proper. The central yolk mass of the yolk cell is surrounded by a cortical cytoplasmic layer comprising two territories: the yolk syncytial layer (YSL) populated by nuclei and underlying deep cells and the EVL and the nuclei-free yolk cytoplasmic layer (YCL). Zebrafish epiboly is driven by a multitude of cell behaviors and physical forces that coordinate epibolic movements of these three cell types (Bruce and Heisenberg, 2020) (C.B. Kimmel et al., 1995) (Kimmel and Law, 1985a) (Kimmel and Law, 1985b). The epiboly onset precedes internalization of mesoderm an endoderm, but upon their formation it involves the expansion and thinning of all three germ layers to eventually enclose the yolk.

The YSL is a crucial source of force generation during the process of epiboly. It forms just before the ZGA, as marginal blastomeres blend into the yolk cytoplasmic layer forming a ring just vegetally to deep cells and the EVL (Chu, 2012; C.B. Kimmel et al., 1995). While in the common syncytial cytoplasm, these nuclei undergo additional mitoses to form two YSL domains. The external YSL (eYSL) is more vegetally located and comprises nuclei not covered by deep cells or EVL, whereas the internal YSL (iYSL) syncytial nuclei are located between the blastoderm and the yolk cell (Chu et al., 2012; C. B. Kimmel et al., 1995). At the initiation of epiboly, the eYSL contracts and the iYSL expands animally, while simultaneously the yolk bulges upwards towards the blastoderm in the process of doming (Solnica-Krezel and Driever, 1994). Doming relies on actomyosin contractility within the YSL, which transmits these pulling forces to the attached EVL as well as deep cells (Bruce and Heisenberg, 2020; Kimmel et al., 1995) (Goonesinghe et al., 2012; Shimizu et al., 2005; Siddiqui et al., 2010; Slanchev et al., 2009). Simultaneously, the yolk microtubule cytoskeleton forms by individual microtubules emanating from microtubule organizing centers (MTOCs) within the external YSL and polymerizing to invade into the YCL and eventually covering the entire exposed yolk. If the integrity or dynamics of these microtubules is disrupted, epiboly is impaired (Fei et al., 2019; Solnica-Krezel and Driever, 1994; Strähle and Jesuthasan, 1993). Nuclei of the external YSL are thought to be transported along this network towards the vegetal pole throughout the process of epiboly (Fei et al., 2019).

Mutational analyses in zebrafish have implicated the atypical cadherin Dachsous1b (Dchs1b) in epiboly (Li-Villarreal et al., 2015). Evolutionarily conserved Dchs proteins consist of a single-pass transmembrane domain, a large extracellular domain with 27 cadherin tandem repeats, and a short intracellular domain (ICD)(Clark et al., 1995). In *Drosophila*, Dachsous is encoded by a single *dachsous* (*ds*) gene, and with its heterophilic binding partner, Fat, in the wing epithelium cells mediates proximal-distal polarization of microtubules that in turn are used to transport PCP signaling components (Harumoto et al., 2010; Matis et al., 2014). In mammals, there are two *dchs* genes, and zebrafish has three: *dchs1a*, *dchs1b*, and *dchs2*. The temporal expression of these three genes suggests both non-overlapping and redundant functions in zebrafish embryogenesis (Li-Villarreal et al., 2015). Accordingly, maternal-zygotic (MZ) *dchs2* mutants exhibit epiboly and mild convergence and extension (C&E) delay without early embryogenesis defects, and double MZ*dchs1b*; MZ*dchs2* mutants display a slightly enhanced epiboly delay phenotype compared to single MZ*dchs1b* mutants (Li-Villarreal et al., 2015). Moreover, *dchs1a* is also expressed during epiboly (Li-Villarreal et al., 2015). MZ*dchs1b* mutants exhibit many defects during early embryogenesis, including abnormal separation of cytoplasm and yolk material in early zygote and irregular cleavages. Mechanistically, Dchs1b promotes microtubule dynamics during early cell cleavages in part by binding via its ICD to Tetratricopeptide repeat protein 28 (Ttc28)(Chen et al., 2018), a protein previously implicated in cell division (Hulpiau and van Roy, 2009; Izumiyama et al., 2012). These studies support a model for Dchs1b function during early cleavages whereby Dchs1b reduces the activity of Ttc28, which limits microtubule dynamics by recruiting it away from the MTOC to the cell membrane (Chen et al., 2018). Whether Dchs employs such mechanisms during epiboly remains to be investigated.

The dynamics of the microtubule cytoskeleton can be influenced by diverse microtubule associated proteins (MAPs) and posttranslational microtubule modifications (Akhmanova and Steinmetz, 2015). K40 acetylation of α-tubulin confers on microtubules resistance to depolymerization and mechanical stress breaks (LeDizet and Piperno, 1986; Piperno et al., 1987; Xu et al., 2017). Since acetylation is a very slow process, only very stable microtubules are acetylated (Howes et al., 2014; Piperno and Fuller, 1985; Szyk et al., 2014). Tyrosination of the C-terminus of α-tubulin can be a reversible or permanent change exerted on tubulin dimers (Berezniuk et al., 2013; Ersfeld et al., 1993; Hallak et al., 1977; Kumar and Flavin, 1981; Paturle-Lafanechère et al., 1991; Rogowski et al., 2010). Detyrosinated microtubules persist longer than tyrosinated ones (Kreitzer et al., 1999) and are protected against depolymerization by Kinesin-13 (Peris et al., 2009; Sirajuddin et al., 2014).

The single Drosophila *ds* regulates the actomyosin cytoskeleton. Accordingly, zebrafish maternal (M)*dchs1b* mutants exhibit defects in processes mediated by actomyosin cytoskeleton - cortical granule exocytosis and cytoplasmic streaming that separates yolk from cytoplasm (Li-Villarreal et al., 2015). During zebrafish epiboly, actin becomes enriched at the EVL and deep cell layer (DEL) margins starting at 50% epiboly and actomyosin flow from vegetal to animal in the yolk cell cortex is observed already at 40% epiboly (Bruce and Heisenberg, 2020). However, whether these actin structures and processes are dependent on *dchs* genes has not yet been addressed.

Here we investigated functions of the three zebrafish *dchs* genes, *dchs1a*, *dchs1b* and *dchs2*, and report that they have partially overlapping functions in the regulation of epiboly progression. By engineering GFP cDNA fragments into the endogenous *dchs1b* locus, we have created Dchs1b-GFP knock-in lines where up to six GFP copies are fused to the C-terminus of Dchs1b ICD. This allowed for the first-time detection of Dchs1b-GFP fusion protein expressed from the endogenous *dchs1b* locus. We observed its localization in both the cell membrane and the cytoplasm of EVL and deep cells during gastrulation. By further studying characteristics of the yolk microtubule network in *dchs* triple loss-of-function mutant embryos, we show that these microtubules are more bundled than in their wild-type (WT) counterparts—a phenotype often associated with defects in epiboly progression (Solnica-Krezel and Driever, 1994). We also show altered yolk cell microtubule polymerization dynamics during epiboly in the MZ*dchs1b* mutant. Moreover, the epiboly progression defects of *dchs1b* mutants were partially suppressed by mutations in the *ttc28* gene. We further demonstrate that MZ*dchs1b* mutants exhibit abnormal organization and dynamics of yolk cell actin cytoskeleton during epiboly. Together, these lines of evidence as well as our transcriptomic analyses support the notion that, like during early embryonic cleavages, Dchs1b has a major while Dchs1a and Dchs2 proteins play supporting roles in promoting microtubule dynamics and organization of both microtubule and actin cytoskeleton to ensure normal epiboly.

## Results

### Zebrafish *dchs1a*, *dchs1b* and *dchs2* genes have overlapping functions during epiboly

The temporal expression of the three zebrafish *dchs* genes suggests both non-overlapping and redundant functions in zebrafish embryogenesis. We have previously reported that *dchs1b* transcripts are heavily maternally loaded and zygotically expressed. By contrast, *dchs1a* and *dchs2* have relatively low maternal expression but higher zygotic expression during gastrulation (Li-Villarreal et al., 2015), although other transcriptomic studies reported higher levels of maternal *dchs1a* expression than that of *dchs1b* (White et al., 2017). Functionally, M*dchs1b* and MZ*dchs1b* mutants are defective in the separation of yolk and cytoplasm, dorsal determinant transport, early cell cleavages and epiboly, indicating that the other two *dchs* genes cannot compensate for its early embryonic functions before ZGA. As loss of *dchs1b* slows epiboly from the shield to (yolk plug closure) YPC stage (Li-Villarreal et al., 2015), we wanted to test if the three *dachsous* genes were functionally redundant during epiboly.

To address this, we did not wish to remove maternal function of *dchs1b* because its loss causes the early defects that could indirectly impair epibolic movements. Instead, in this analysis we tested if zygotic functions of the *dchs1b* gene were redundant with maternal and zygotic functions of *dchs1a* and *dchs2*, by comparing single Z*dchs1b^-/-^* mutants to triple MZ*dchs1a^-/-^*; Z*dchs1b^-/-^*; MZ*dchs2^-/-^* mutants, henceforth referred to as *dchs* triple mutants. The MZ*dchs1a^-/-^*; *Zdchs1b^+/-^*; MZ*dchs2^-/-^* will be referred to as *dchs1a;2* double mutants. We used confocal microscopy with high temporal resolution, to examine embryos in which nuclei and membranes were labeled by injecting synthetic mRNAs coding for fluorescently tagged Histone H2B-RFP and mCherry-CAAX, respectively (Figure 1). Epiboly defects are often described in both overall progression of deep cells and EVL towards the vegetal pole as well as the separation of the most superficial EVL, from the deep cells, which are lagging behind EVL in their epibolic movement (Figure 1B schema). Compared to wild-type (WT) controls, Z*dchs1b* mutants exhibited very mild epiboly progression defects and a slight increase in separation between the EVL and DEL; these defects were much stronger in *dchs1a;2* double and *dchs* triple mutants (Figure 1A,C,D; Supplement Figure 1). Two MZ*dchs1a^-/-^*; MZ*dchs2^-/-^* compound mutant combinations, one with Z*dchs1b^+/-^*and the other with Z*dchs1b^-/-^*, showed more severe and significant epiboly delay and larger separation between the EVL and DEL. However, the *dchs* triple mutants exhibited the most severe delay in epiboly and largest and most sustained separation between the EVL and DEL amongst the genotypes analyzed (Figure 1A,C,D; Supplement Figure 1). Further, more severely affected *dchs* triple mutant embryos had jagged DEL margins, contrasting the smooth epibolic front of deep cells in WT (Figure 1A). Additionally, two out of six embryos failed to complete epiboly even after ten hours post-oblong stage, while their WT counterparts completed yolk plug closure by 6.8 hours post-oblong stage (Supplement Figure 1). Together, these data show that triple *dchs* mutants exhibit the most penetrant and consistent epiboly delay and DEL-EVL separation. Importantly, this finding, paired with the milder epiboly defects in Z*dchs1b^-/-^*, suggests these three *dchs* genes have semi-overlapping and redundant functions during epiboly.

**Figure 1.**
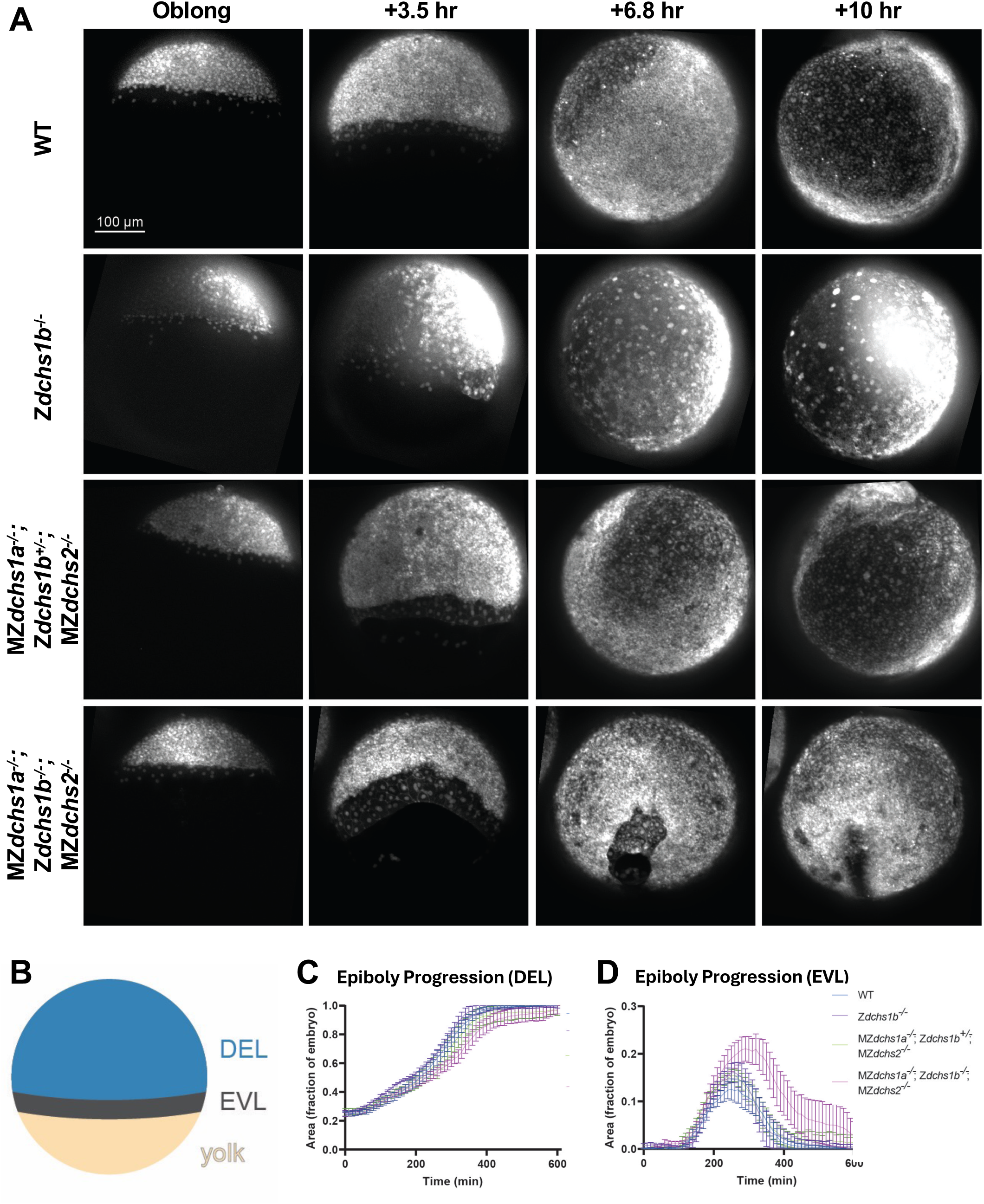
Epiboly progression and EVL-DEL separation defects in Z*dchs1b* and *dchs* triple mutants. A. Representative confocal images of WT, Z*dchs1b^-/-^*, MZ*dchs1a^-/-^*; Z*dchs1b^+/-^*; MZ*dchs2^-/-^*, and MZ*dchs1a^-/-^*; Z*dchs1b^-/-^;* MZ*dchs2^-/-^*, with nuclei labeled with H2B-RFP and membranes labeled with mCherry-CAAX beginning at the oblong stage, then 3.5, 6.8, and 10-hours post-oblong. B. Diagram of a mid-epiboly stage embryo with the deep cell layer (DEL, blue), enveloping layer (EVL, dark gray), and yolk (yellow) labeled. Blue area corresponds to data presented in C, and the dark gray area corresponds to data presented in D. C. Quantification of DEL progression towards the vegetal pole during epiboly. Error bars indicate SEM. WT N=7 embryos. Z*dchs1b^-/-^* N=6 embryos. MZ*dchs1a^-/-^*; Z*dchs1b^+/-^*; MZ*dchs2^-/-^* N=6 embryos. MZ*dchs1a^-/-^*; Z*dchs1b^-/-^*; MZ*dchs2^-/-^* N=6 embryos. Statistical significance calculated with one-way ANOVA Friedman test corrected for multiple comparisons. WT vs. Z*dchs1b^-/-^*, p<0.01; WT vs. MZ*dchs1a^-/-^;* Z*dchs1b^+/-^*; MZ*dchs2^-/-^*, ns; WT vs. MZ*dchs1a^-/-^;* Z*dchs1b^-/-^;* MZ*dchs2^-/-^*, p<0.0001; Z*dchs1b^-/-^* vs. MZ*dchs1a;* Z*dchs1b^+/-^*; MZ*dchs2^-/-^*, p<0.0001; Z*dchs1b^-/-^*vs. MZ*dchs1a^-/-^;* Z*dchs1b^-/-^*; MZ*dchs2^-/-^*, p<0.0001; MZ*dchs1a;* Z*dchs1b^+/-^*; MZ*dchs2^-/-^*vs. MZ*dchs1a^-/-^;* Z*dchs1b^-/-^*; MZ*dchs2^-/-^*, p<0.0001. D. Quantification of the separation between the EVL and DEL during epiboly. Error bars indicate SEM. WT N=7 embryos. Z*dchs1b^-/-^*N=6 embryos. MZ*dchs1a^-/-^*; Z*dchs1b^+/-^*; MZ*dchs2^-/-^* N=6 embryos. MZ*dchs1a^-/-^*; Z*dchs1b^-/-^*; MZ*dchs2^-/-^* N=6 embryos. Statistical significance calculated with one-way ANOVA Friedman test corrected for multiple comparisons. WT vs. Z*dchs1b^-/-^*, p<0.001; WT vs. MZ*dchs1a^-/-^;* Z*dchs1b^+/-^*; MZ*dchs2^-/-^*, p<0.0001; WT vs. MZ*dchs1a^-/-^;* Z*dchs1b^-/-^;* MZ*dchs2^-/-^*, p<0.0001; Z*dchs1b^-/-^* vs. MZ*dchs1a;* Z*dchs1b^+/-^*; MZ*dchs2^-/-^*, ns; Z*dchs1b^-/-^* vs. MZ*dchs1a^-/-^;* Z*dchs1b^-/-^*; MZ*dchs2^-/-^*, p<0.0001; MZ*dchs1a;* Z*dchs1b^+/-^*; MZ*dchs2^-/-^* vs. MZ*dchs1a^-/-^;* Z*dchs1b^-/-^*; MZ*dchs2^-/-^*, p<0.001.

### Endogenous Dchs1b-6sfGFP fusion protein localizes to cell membrane and cytoplasm of deep and enveloping layer cells during epiboly

As a single-pass transmembrane protein, Ds is localized to cell membranes in *Drosophila* epithelia. Antibody staining indicated that the murine Dchs1 is ubiquitously expressed in cell membranes of mouse embryos and in cultured human respiratory epithelial cells DCHS1 is localized to the base of ciliary apparatus (White et al., 2017). We previously reported that the zebrafish Dchs1b-sfGFP C-terminal fusion protein overexpressed in early zebrafish embryos was detected at cell membranes and cytoplasm of early blastomeres before the epiboly onset (Chen et al., 2018).

To characterize the intracellular distribution of the endogenous Dchs1b protein in live embryos undergoing epiboly, we edited the endogenous *dchs1b* gene to insert two or six copies of sfGFP into the *dchs1b* stop codon to generate a Dchs1b fusion protein with a fluorescent tag at its C-terminus (Figure 2A,B; Materials and Methods). The resulting knock-in homozygous embryos generated by homozygous parents did not show any overt defects in yolk-cytoplasm separation or early cleavages observed in MZ*dchs1b* nonsense mutants (Chen et al., 2018; Li-Villarreal et al., 2015), or strong epiboly defects (data not shown). Therefore, we concluded that this tag does not interfere with WT *dchs1b* function.

**Figure 2.**
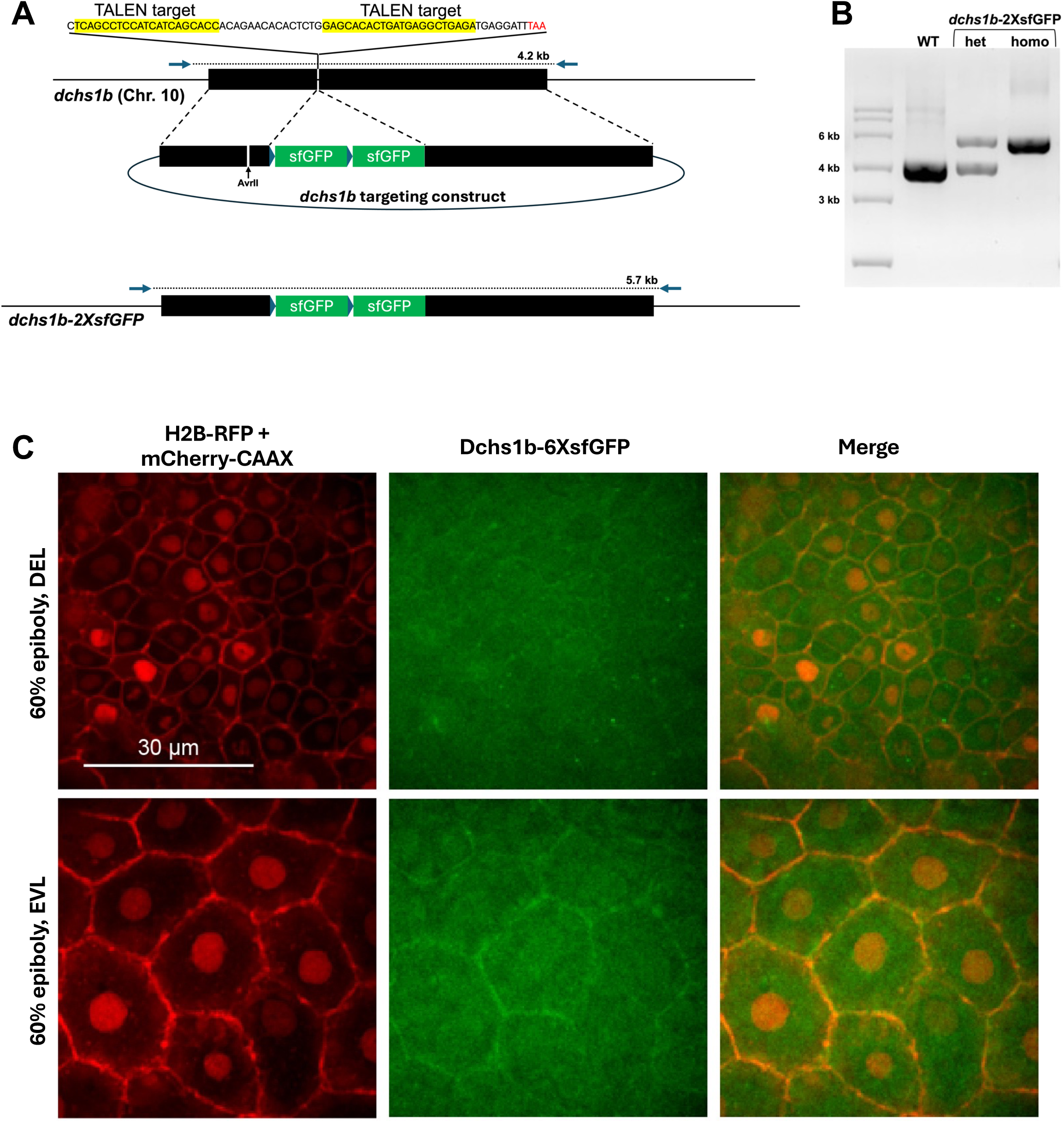
Endogenous expression of Dchs1b in the membrane and cytoplasm during epiboly. A. Schema of generating *dchs1b-2xsfGFP* line. B. Agarose electrophoresis gel showing genotyping for the endogenous dchs1b locus without and with *2xsfGFP* insertion. C. Confocal images at 60% epiboly with nuclei and membranes labeled with H2B-RFP (red) and mCherry-CAAX (red) along with endogenous expression of Dchs1b tagged with six copies of sfGFP (green) in both the EVL and DEL.

Using confocal microscopy imaging of MZ*dchs1b-2xsfGFP* embryos, we could not detect the fusion protein at any stages of early embryogenesis. By contrast, examination of MZ*dchs1b-6xsfGFP* embryos co-expressing H2B-RFP and mCherry-CAAX proteins at 60% epiboly, showed accumulation of the fusion protein at EVL cell membranes, weak membrane localization in deep cells and a cytoplasmic signal (Figure 2C). These observations indicate that Dchs1b-6xsfGFP localizes to cell membranes of EVL and deep cells during epiboly and suggest the presence of its C-terminal fragments in the cytoplasm.

### *dchs* deficiency leads to structural defects in the dynamic yolk cell microtubule array

Perturbations in cell adhesion and/or the yolk cytoskeleton can cause epiboly defects (Bruce and Heisenberg, 2020). Dachsous is involved in regulating microtubules in *Drosophila* (Harumoto et al., 2010; Matis et al., 2014). Our work showed that during early embryonic cleavages microtubules in MZ*dchs1b* mutants are less dynamic (Chen et al., 2018), while during epiboly the yolk cell microtubules are bundled, and the elaborate microtubule meshwork is discontinuous (Li-Villarreal et al., 2015). Hence, we analyzed the yolk microtubule array in *dchs* triple mutants during epiboly in more detail. Consistent with previous findings in MZ*dchs1b* mutant embryos (Li-Villarreal et al., 2015), *dchs* triple mutants displayed yolk microtubule bundling and associated “naked” yolk regions without microtubules, with an average yolk area lacking microtubules of 12.7% compared to just 0.6% in WT (Figure 3A,B). In the *dchs* double mutant embryos the yolk area devoid of microtubules comprised on average 11.1%, and while this was not significantly different from *dchs* triple mutants, the bundling phenotype was less penetrant in *dchs* double mutant than in triple mutant embryos (37.9% double versus 59.4% of triple *dchs* mutant embryos) (Supplement Figure 2). These data, paired with more severe epiboly progression defects in *dchs* triple mutants, indicate triple *dchs* mutant embryos manifest the most severe and penetrant yolk microtubule and epiboly progression defects. Therefore, more detailed analyses of the yolk cell microtubule array were conducted in this genetic background.

**Figure 3.**
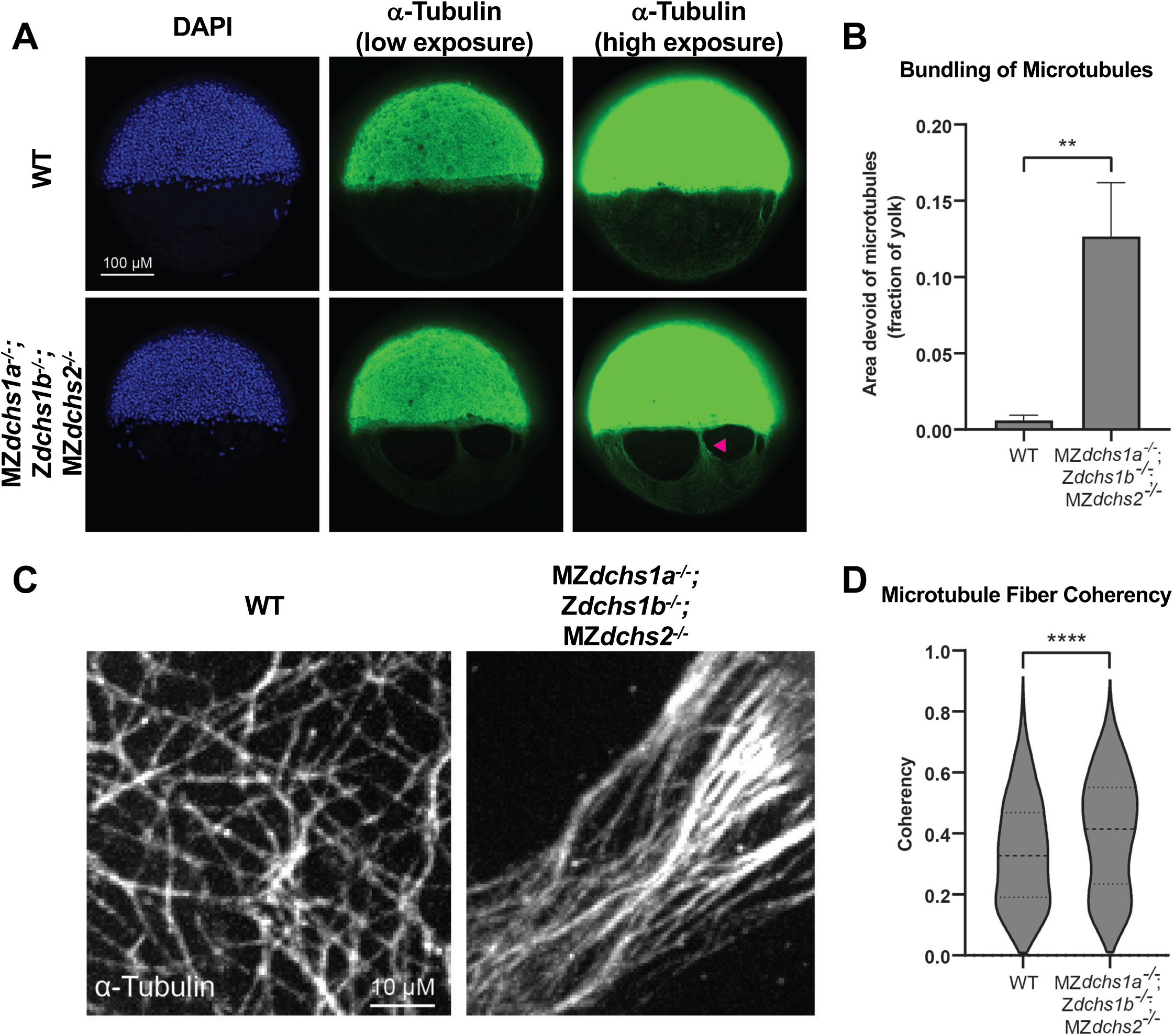
Bundled yolk microtubule array structures in *dchs* triple mutants. A. Confocal images at 50% epiboly of WT and *dchs* triple mutants stained with DAPI (blue) and for DM1α (microtubules, green). Arrowhead indicates microtubule bundling. B. Quantification of yolk microtubule bundling through measurement of bare yolk areas lacking microtubules at 50% epiboly. **p<0.01. WT N=25 embryos, MZ*dchs1a^-/-^*; Z*dchs1b^-/-^*; MZ*dchs2^-/-^* N=32 embryos. C. Confocal images of yolk microtubule meshwork marked with DM1α, just vegetal of YSL of WT and *dchs* triple mutants at 50% epiboly. D. Quantification of yolk microtubule alignment coherency. ****p<0.0001. WT N=25 embryos, MZ*dchs1a^-/-^*; Z*dchs1b^-/-^*; MZ*dchs2^-/-^* N=32 embryos.

To better understand differences in the microtubule cytoskeleton organization, we analyzed higher magnification images of yolk microtubules for their overall alignment. In WT embryos, the yolk microtubules form a complex, basket-like meshwork to evenly cover areas of the yolk (Figure 3C) (Solnica-Krezel and Driever, 1994; Strähle and Jesuthasan, 1993). However, in *dchs* triple mutant embryos, microtubule fibers were generally more aligned with one another within the network (Figure 3C). Quantitatively, we analyzed the coherency of microtubules located just vegetally to the YSL. The overall coherency of these sample areas was significantly higher in *dchs* triple mutants (0.4015±0.0053, 95% CI) compared to WT (0.3388±0.0054, 95% CI), indicating microtubule fibers in *dchs* triple mutants are more aligned (Figure 3D). These studies revealed that there were overall structural defects in the yolk microtubule cytoskeleton in *dchs* triple mutants but determining how these defects arise requires analysis of microtubule dynamics, microtubule binding proteins, and/or microtubule post-translational modifications.

### *dchs* deficiency leads to abnormal microtubule polymerization dynamics

To measure dynamics at the plus-ends of microtubules, we injected WT and *dchs* triple mutants at the one cell stage with *EB3-GFP* synthetic mRNA, which encodes a plus-end binding protein often used to monitor microtubule polymerization (Stepanova, 2003). We collected time-lapse movies at 4-4.5 hours post fertilization (hpf), or dome stage, when the yolk microtubule network is being established. By viewing maximum time projections of these time-lapse movies, we monitored the paths of EB3 track histories. Compared to WT, the polymerization of yolk microtubules in *dchs* triple mutant appeared spatially uneven with certain regions containing disorganized polymerization activity (Figure 4A, magenta arrowhead). Microtubule polymerization tracks showed a statistically significant increase of total duration in *dchs* triple mutants compared to WT (Figure 4B). The distance between the first and last detected EB3-GFP comet of a single microtubule polymerization event, or track displacement, was also significantly increased in *dchs* triple mutants compared to WT (Figure 4C). Average microtubule track speed trended towards an overall increase compared to WT, but this was not significant (Figure 4D). Analysis of MZ*dchs* triple mutants compared to wild type yielded similar results, significant increase in track duration and displacement and a trend towards faster track speeds (Supplement Figure 3). Together, these data indicate that *dchs* genes play a role in limiting microtubule polymerization and promoting dynamics during epiboly.

**Figure 4.**
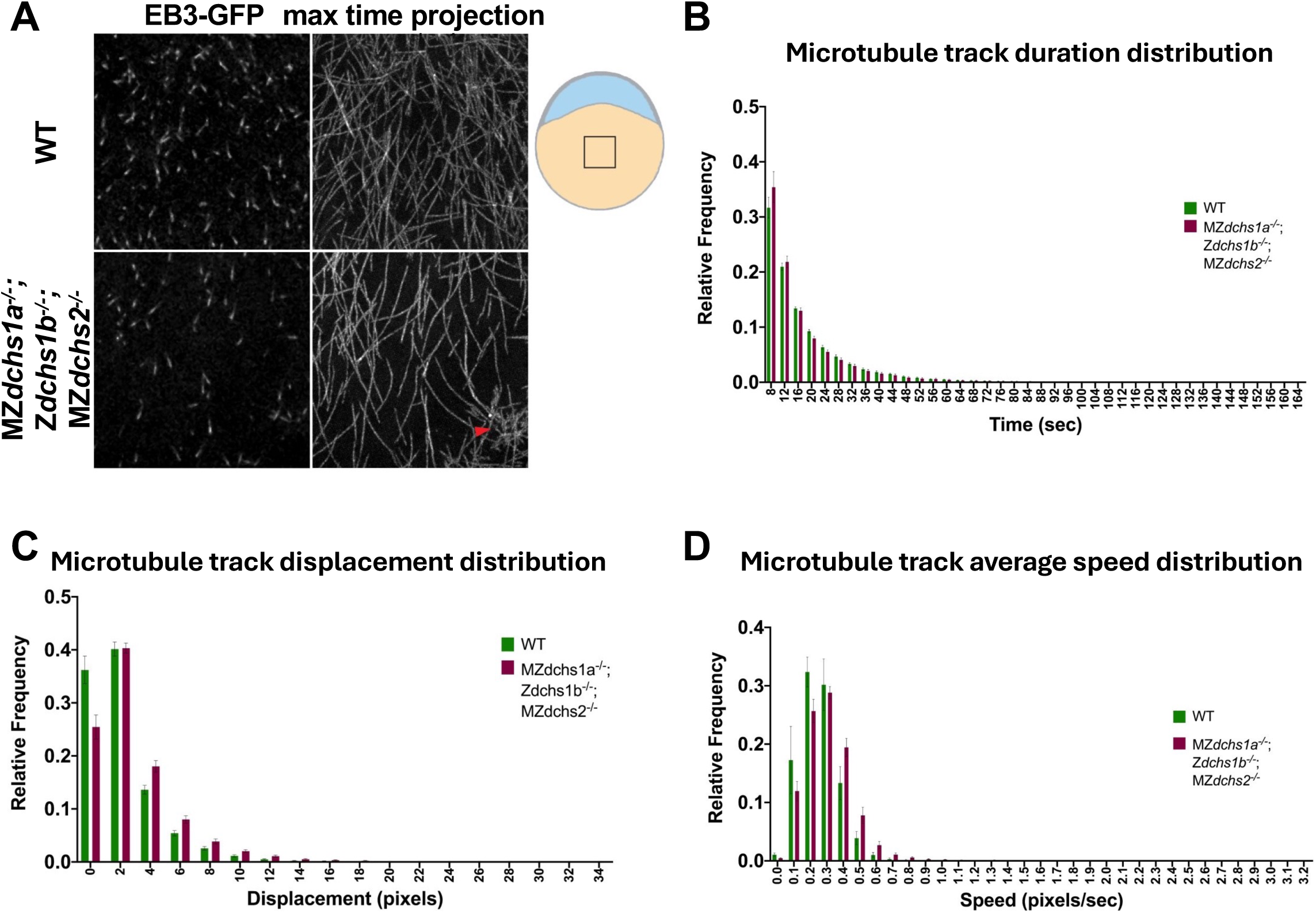
Yolk microtubule polymerization in *dchs* triple mutants. A. Sample confocal images (left) and maximum time projections of time lapses (right) of WT and *dchs* triple mutants labeled with EB3-GFP. Arrowhead indicates disorganized area of polymerization. Embryo schema on the right with the box illustrating YCL imaging region. B. Frequency distribution of EB3 track duration. Error bars indicate SEM. WT N=9 embryos, n=26,573 tracks. MZ*dchs1a^-/-^*; Z*dchs1b^-/-^*; MZ*dchs2^-/-^* N=18, embryos n=29,668. Wilcoxon matched-pairs rank test, *p<0.05. C. Frequency distribution of EB3 track displacement. Error bars indicate SEM. WT N=9 embryos, n=26,573 tracks. MZ*dchs1a^-/-^*; Z*dchs1b^-/-^*; MZ*dchs2^-/-^* N=18, embryos n=29,668. Wilcoxon matched-pairs rank test, **p<0.01. D. Frequency distribution of EB3 track average speed. Error bars indicate SEM. WT N=9 embryos, n=26,573 tracks. MZ*dchs1a^-/-^*; Z*dchs1b^-/-^*; MZ*dchs2^-/-^* N=18, embryos n=29,668. Wilcoxon matched-pairs rank test, ns, *p<0.05.

### Microtubule bundles are generally associated with increased acetylation, but not tyrosination

Certain types of posttranslational microtubule modifications are considered markers of and can even confer stability, such as K40 acetylation and detyrosination (Akhmanova and Steinmetz, 2015). To test whether such modifications are altered in *dchs* triple mutants, we performed immunohistochemistry at 60% epiboly. In both WT and triple *dchs* mutant embryos, about 77-78% of the yolk microtubules exhibited similar levels of tyrosinated tubulin (Figure 5A,B). Acetylated tubulin was detectable at mitotic spindles, midbodies, as well as yolk microtubule bundles in WT and *dchs* triple mutant embryos (Figure 5C). There were no clear differences in acetylated microtubules between WT and *dchs* triple mutant embryos. Because bundles of microtubules in both WT and *dchs* triple mutant embryos were associated with acetylated tubulin signal, it was not feasible to discern whether microtubule bundling in general lead to acetylation, or if acetylation leads to increased bundling. However, as noted above, microtubule bundles were rare in WT and common in *dchs* triple mutants (Figure 3B). Together, these data suggest yolk microtubule bundles in general are associated with acetylated tubulin during epiboly.

**Figure 5.**
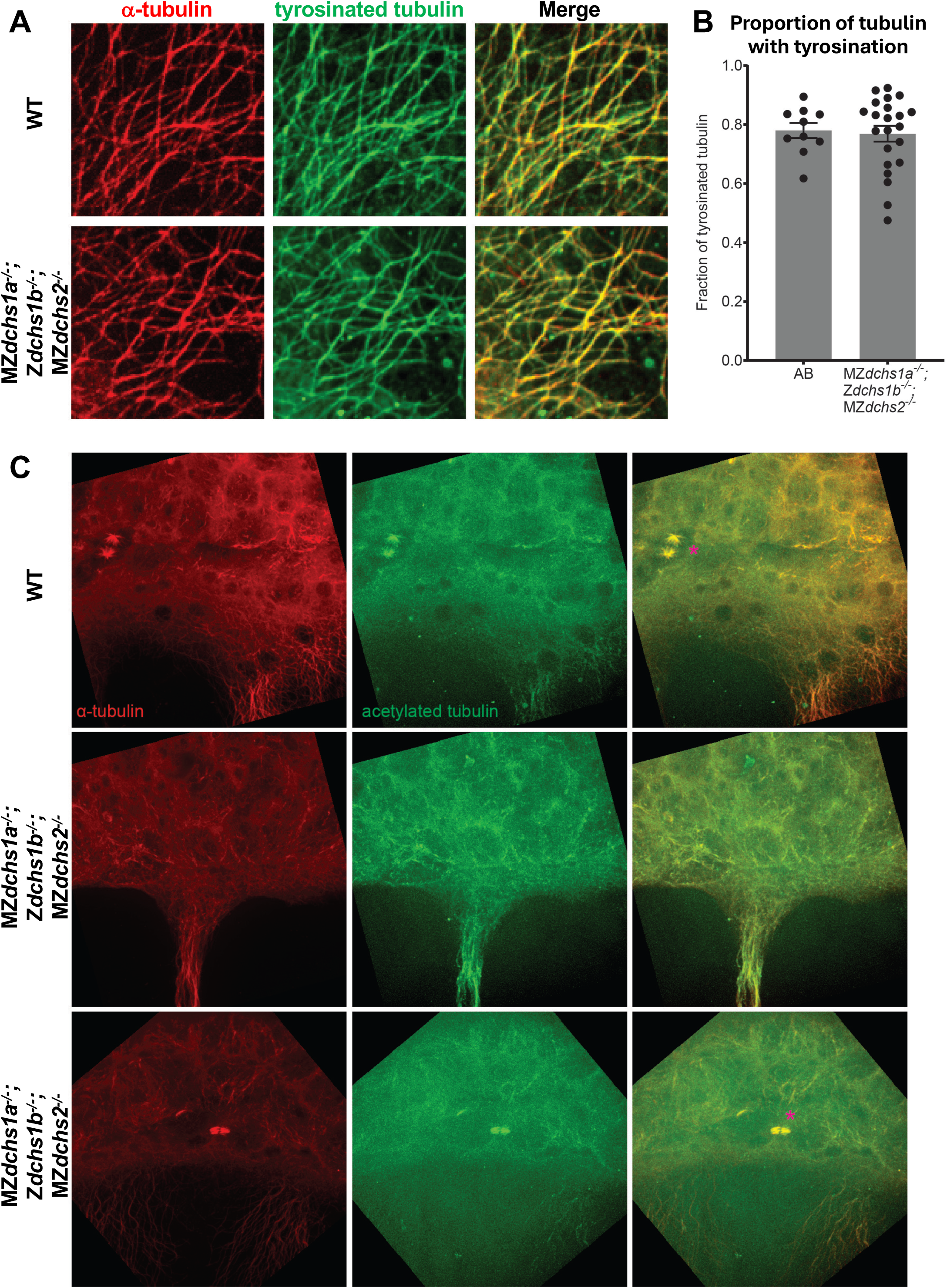
Tyrosinated and acetylated tubulin modifications in *dchs* triple mutants. A. Confocal images of yolk microtubules (DM1α, red) and tyrosinated tubulin (green) in both WT and *dchs* triple mutants. B. Quantification of tyrosinated tubulin staining associated with yolk microtubules. Error bars indicate SEM. C. Confocal images of yolk microtubules (DM1α, red) and acetylated tubulin (green) in both WT and *dchs* triple mutants. Asterisks in merged channels indicate microtubule spindles.

### Dachsous promotes epiboly by limiting Ttc28 function

We have previously reported that Dchs1b interacts with Ttc28 and AuroraB to control microtubule dynamics during cleavage stages. Specifically, Dchs1b promotes microtubule dynamics by regulating intracellular distribution of Ttc28, which limits microtubule dynamics. Interestingly, reduced yolk cell microtubule dynamics phenotype during cleavage stages of M*dchs1b* mutants was suppressed in M*dchs1b*; M*ttc28* double mutants (Chen et al., 2018). MZ*ttc28* mutants do not present with overt epiboly defects (Chen et al., 2018). To test whether Dchs1b also interacts with Ttc28 during epiboly we analyzed epiboly progression and separation of EVL and deep cells in single MZ*dchs1b* and MZ*ttc28* mutants, MZ*dchs1b*; MZ*ttc28* double mutants as well as their WT siblings (Figure 6A,B). Separation of EVL and deep cells was comparable between MZ*ttc28-/-* and WT. As expected, the EVL-DEL separation was significantly increased in MZ*dchs1b* mutants but notably it was normalized in MZ*dchs1b*; MZ*ttc28* double mutants (Figure 6B). These genetic interactions studies support the model that Dsch1b functionally interacts with Ttc28 by negatively regulating its activity in many morphogenetic processes driven by microtubules.

**Figure 6:**
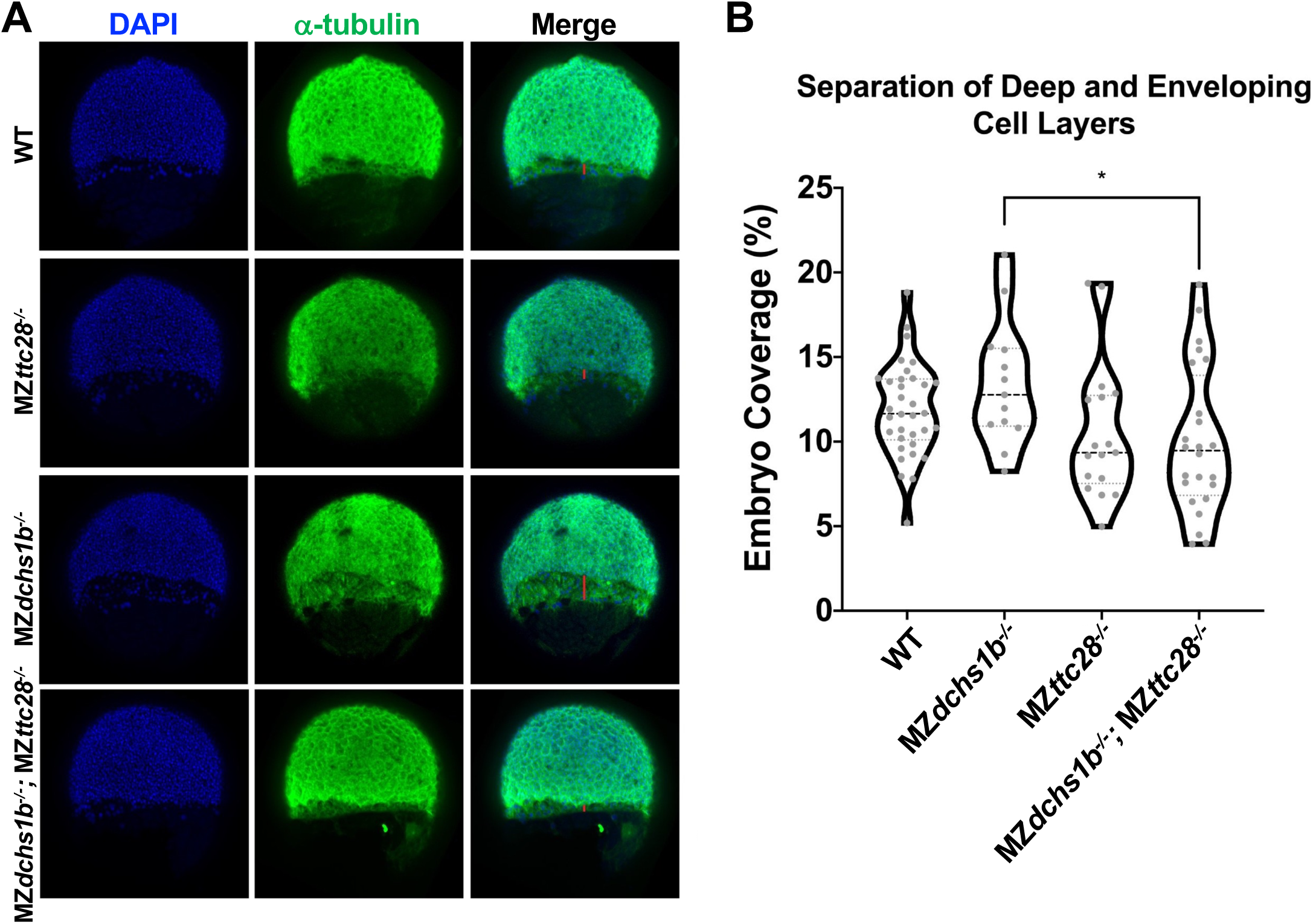
Interaction between *dchs1b* and *ttc28* genes in epiboly progression. A. Representative confocal images (from the top to bottom row) of WT, MZ*ttc28^-/-^*, MZ*dchs1b^-/-^,* and MZ*dchs1b^-/-^*; MZ*ttc28^-/-^* with nuclei labeled with DAPI and yolk microtubules with DM1α (green) at 60% epiboly. B. Quantification of the separation between the EVL and DEL during epiboly in embryos of the above genotypes. The following number of embryos from four different clutches were analyzed: WT, n=33; MZ*ttc28^-/-^*, n=17; MZ*dchs1b^-/-^*, n=37; MZ*dchs1b^-/-^*;MZ*ttc28^-/-^*, n=23.

### Dachsous deficiency impairs the dynamic yolk actin cytoskeleton

Actin cytoskeleton is essential for zebrafish epiboly. Actin becomes enriched at the EVL and DEL margins starting at 50% epiboly and actomyosin flow from vegetal to animal in the yolk cell cortex is initiated at 40% epiboly (Bruce and Heisenberg, 2020). As M*dchs1b* mutants exhibit defects in actin-dependent processes such as cortical granule exocytosis and separation of yolk and cytoplasm in the activated zebrafish egg (Li-Villarreal et al., 2015), we asked whether Dchs function is required for the dynamic actin cytoskeleton organization during epiboly. We introduced *Tg(μ-actin:utrophin-GFP)* transgene into the *dchs1b* mutant background and analyzed reorganization of the actin cytoskeleton over the course of epiboly (Movie 1). We observed that actin became enriched at the EVL margin starting at 50% epiboly in both WT and mutant embryos (Figure 7A,B). WT embryos showed a uniform actin meshwork on the yolk cell that flowed towards the EVL margin (Figure 7A; Movie 1). By contrast, in MZ*dchs1b* mutant gastrula the actin meshwork was not continuous with clear gaps (Figure 7B) reminiscent of the gaps observed in the yolk cell microtubule network (Figure 3). Whereas the actin network exhibited the vegetal to animal flow towards the EVL margin in WT, this flow was not uniform in MZ*dchs1b* with cables of actin fibers alternating with actin free regions at the EVL margin. Together, these analyses indicate that Dchs1b is required for the dynamic organization of actin cytoskeleton during epiboly in addition to its role in the microtubule network.

**Figure 7:**
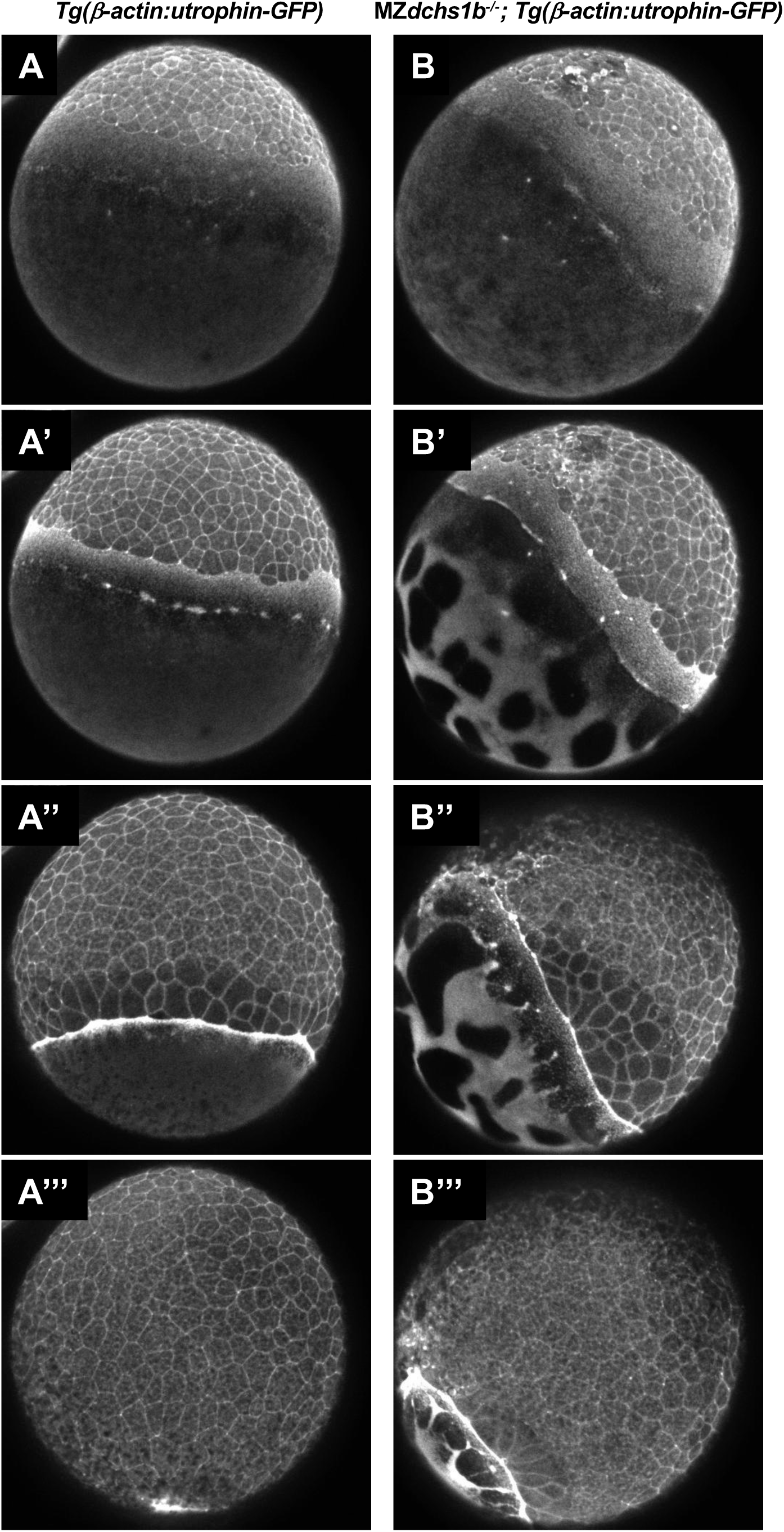
Images of the actin cytoskeleton organization in WT embryos (A, A’,A’’, A’’’) and MZ*dchs1b* mutants (B, B’, B’’, B’’’) harboring *Tg(μ-actin:utrophin-GFP)* transgene that were captured from Supplemental Movie 1 at 30% epiboly (based on WT embryo epiboly progression) (A,B), 40% epiboly (A’,B’); 60% epiboly (A’’, B’’), and 90% epiboly (A’’’, B’’’).

### Altered transcriptome of *dachsous*-deficient gastrulae

We wished to understand if *dachsous* deficiency affects the transcriptome of zebrafish gastrula. We isolated RNA and performed bulk RNA-seq analysis at 10 hpf of WT (AB*) gastrulae and MZ*dchs* (MZ*dchs1a*; MZ*dchs1b*; MZ*dchs2*) mutants, which we separated into two phenotypic classes: mild epiboly delay and severe, which failed to complete epiboly (Figure 8A). A small fraction of triple mutant embryos died before 10 hpf and was not included in the analysis.

**Figure 8:**
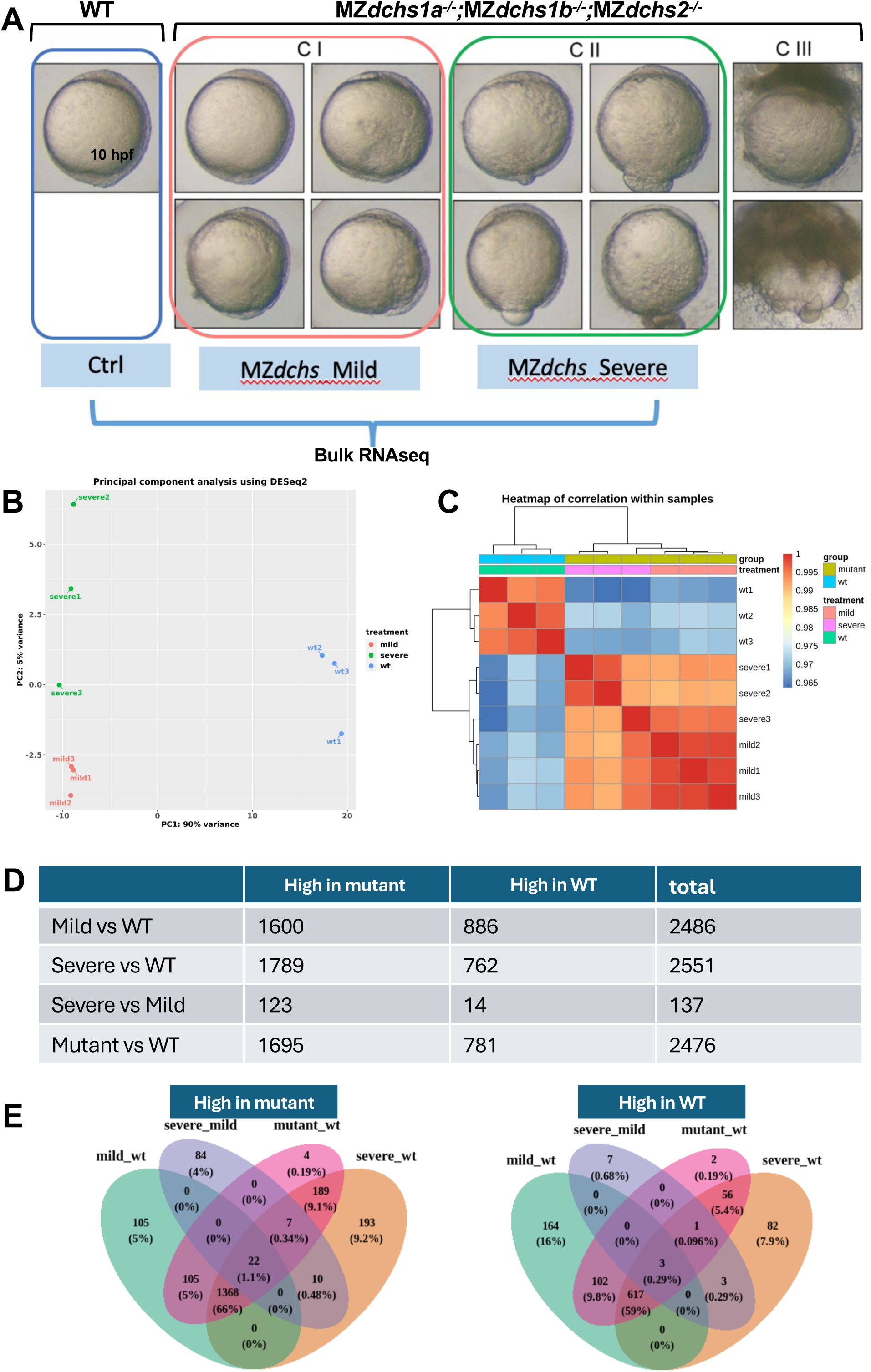
Altered transcriptome in MZ*dchs1a,b,2* triple mutants at the end of epiboly (10hpf). A. Representative dissecting microscope images of WT and the three phenotypic classes of MZ*dchs1a,b,2* triple mutant embryos. Only classes CI and CII were collected for bulk RNA sequencing analysis. B. Principal component analysis using RNA sequencing data. Percent variability within each principal component is listed on each axis. C. Heatmap of correlation of 3 independent RNA samples of WT, mild and severely affected MZ*dchs1a,b,2* triple mutant embryos. The color represents the strength and direction of correlation. D. Table presenting the number of differentially expressed genes in WT and the mild and severely affected MZ*dchs1a,b,2* triple mutant embryos (high in mutant or in WT). E. Venn diagrams representing the overlap and differences in gene expression between genes upregulated (left) or downregulated (right) in the mild and severely affected MZ*dchs1a,b,2* triple mutant embryos relative to WT.

Bioinformatic analysis identified 2,476 differentially expressed genes (DEGs) between WT gastrulae and MZ*dchs* mutants with strong phenotype and 2,383 between WT MZ*dchs* mutants with mild phenotype. By several measures, transcriptomes of MZ*dchs* mutants with mild and severe phenotypes appeared to be very similar (Figure 8B,C,D,E). Mild and severe mutants shared all phenotypes except epiboly completion by 10hpf. Accordingly, principal component analysis of the transcriptomes of WT, mild and strong MZ*dchs* mutants showed that both mutant classes are similarly distanced from WT along the PC1 component, which accounted for 92% of variance between WT and the two mutant classes. Moreover, the mild and severe mutants overlapped on PC2 with a spread over the samples, accounting for only 5% of the variance (Figure 8B). Similarly, the transcriptome correlation heatmap showed clear differences in gene expression between WT vs. mutant datasets but a high similarity between the mutants with mild and severe phenotypes (Figure 8C). Finally, comparison of the genes with two-fold or greater up or downregulation and an adjusted p-value of less than 0.05, showed a ∼1:4:1 ratio (481:1902:572) of DEGs found exclusively in the transcriptome of mutants with mild phenotype, DEGs shared by both mild and severe phenotype to DEGs found exclusively in the mutants exhibiting severe phenotype (not shown). We interpret these analyses to mean that the observed differences in the epiboly delays in the two mutant classes are not likely due to large differences in their transcriptomes.

FishEnricher pathway analysis of 781 genes downregulated in *dachsous* mutants indicated “cortical actin cytoskeleton” and “intermediate filament” as Cellular Components (Supplement Figure 4A,B,C,D). Intriguingly “mitochondrial fission” and “mitochondrial fusion” were indicated as GO Biological Process and Phenotype, while “Chorion” as Anatomy and Phenotype. FishEnrichr analysis of the 1,697 genes upregulated in the mutants in the Biological Process category pointed to “Striated muscle”, “myofibril assembly”, “cardiac muscle development”, including genes and Positive regulation of “cell-substrate adhesions” with several collagen-encoding genes in this DEG category (*col9a3, col3a2, col18a1, col2a1b*) (Supplement Figure 4E). Altogether, these transcriptomic analyses indicate high similarity between mutants with various expressivity of the epiboly defect. Moreover, the differentially expressed genes point to dysregulated actin and intermediate cytoskeleton, as well as cell-cell and cell matrix adhesion.

## Discussion

Here, we have investigated the functional redundancies between the three zebrafish *dachsous* homologs, *dchs1a*, *dchs1b*, and *dchs2* and the mechanisms by which they contribute to epiboly during zebrafish gastrulation. Our previous study implicated maternally deposited *dchs1b* transcripts in numerous processes during early embryogenesis before ZGA, including egg activation and early cleavages, and maternally and zygotically expressed *dchs1b* in the process of epiboly. The epiboly defects of MZ*dchs1b* mutants are enhanced by inactivation of *dchs2,* (Li-Villarreal et al., 2015). However, the potential functional redundancies between the three *dchs* genes have not been studied. Utilizing the nonsense *dchs1a^sa15468^* allele (Kettleborough et al., 2013), we observed that neither single zygotic nor MZ*dchs1a* mutants exhibited overt embryogenesis defects. Interestingly, MZ*dchs1a*; Z*dchs1b* exhibit stronger epiboly progression defects than single Z*dchs1b* mutants, with these defects further exacerbated in MZ*dchs1a*; Z*dchs1b*; MZ*dchs2*, supporting partially redundant function of these three genes (Figure 1). Our zebrafish studies are consistent with the previous reports in the mouse, where the murine *Dchs1* and *Dchs2* paralogues act in partially redundant fashion to control the number of nephron progenitors (Bagherie-Lachidan et al., 2015).

The epiboly defects of MZ*dchs1a*; Z*dchs1b*; MZ*dchs2* triple mutants likely do not reflect a complete loss of Dachsous function, as MZ*dchs1b* exhibit stronger epiboly impairment than Z*dchs1b* mutants. However, the removal of maternal *dch1b* function impairs early cleavages and separation of yolk and cytoplasm that can indirectly delay epiboly. Our previous qRT-PCR analyses of the *dchs* gene expression are consistent with *dchs1b* being most highly expressed maternally. Contrary to this, bulk transcriptomic studies indicated higher maternal expression of *dchs1a* compared to *dchs1b* and negligible *dchs2* maternal expression (White et al., 2017). The two studies agree about comparable expression levels of *dchs1a* and *dchs1b* during epiboly. Whereas qRT-PCR studies detected *dchs2* transcripts at levels like those of *dch1a* and *dchs1b*, bulk transcriptomic analyses fail to detect *dchs2* during gastrulation. Our RNA-seq transcriptomic analyses at the completion of epiboly, detected *dchs1b* transcripts but background levels of *dchs1a* and *dchs2*. We provided further insight into the *dchs1b* gene expression by inserting into the *dchs1b* stop codon two or six copies of sfGFP. The resulting Dchs1b-6xsfGFP fusion protein was detected during epiboly stages corroborating *dchs1b* expression (Figure 2C).

Both *dchs1b-2xsfGFP* and *dchs1b-6xsfGFP* knock-in homozygous embryos generated by homozygous parents did not show any overt defects in yolk-cytoplasm separation or early cleavages observed in MZ*dchs1b* nonsense mutants (Chen et al., 2018; Li-Villarreal et al., 2015), or strong epiboly defects arguing that the Dchs1b protein fused at its C-terminus with fluorescent tags is functional. We observed the fusion protein at cell membranes of enveloping cells and deep cells during epiboly, consistent with the intracellular distribution of this cadherin in *Drosophila* and in mouse. Significant levels of GFP signal in the cytoplasm of *dchs1b-6xsfGFP* knock-in homozygous embryos could be due to expression of Dchs1b intracellular domain in the cytoplasm or could be due to nonspecific degradation of the fusion protein releasing the GFP portion.

Epiboly defects are hallmark of the *dachsous* triple mutants in zebrafish (Figure 1) as previously reported for MZ*dchs1b*; MZ*dchs2* double mutants (Li-Villarreal et al., 2015). These epiboly defects were correlated with abnormally bundled microtubule organization in the yolk syncytial layer (Li-Villarreal et al., 2015). Such bundled microtubule network and epiboly delays have been previously observed in zebrafish embryos treated with Taxol, a small molecule that stabilizes microtubules (Solnica-Krezel and Driever, 1994), suggesting that Dachsous may regulate microtubule dynamics. Consistent with this notion, our previous studies of M*dchs1b* mutants during early cleavages showed that yolk cell microtubules are less dynamic. Moreover, Dchs1b promotes microtubule dynamics during early cleavages in part by binding via its ICD to Tetratricopeptide repeat protein 28 (Ttc28)(Chen et al., 2018), which had been implicated in mammalian cell division (Hulpiau and van Roy, 2009; Izumiyama et al., 2012). These studies support a model whereby during early cleavages Dchs1b reduces the activity of Ttc28, which limits microtubule dynamics by recruiting Ttc28 away from the MTOC to the cell membrane (Chen et al., 2018; Li-Villarreal et al., 2015). We provided several lines of evidence that Dachsous cadherins employ similar mechanisms to regulate microtubule cytoskeleton and drive epiboly. First, yolk microtubules of MZ*dchs1a*; Z*dchs1b*; MZ*dchs2* triple mutants are bundled (Figure 3A,B), and the bundled microtubules are strongly aligned compared to yolk microtubules of WT gastrulae (Figure 3C,D). Second, live imaging of yolk cell microtubules with plus-end binding EB3-GFP protein during epiboly showed that microtubules in *dchs* triple mutants exhibit longer displacement and a trend towards faster track speeds (Figure 4), consistent with *dchs* genes limiting microtubule polymerization and promoting dynamics, as shown during early cleavage stages (Chen et al., 2018). Finally, supporting the notion that Dachsous promotes epiboly progression by negatively regulating Ttc28, the increased EVL-DEL separation in MZ*dchs1b* mutants was partially suppressed in MZ*dchs1b*; MZ*ttc28* double mutants (Figure 6). Further, Dachsous regulates microtubules via similar mechanisms at different developmental stages and different morphogenetic processes in zebrafish.

In addition to microtubules, the dynamic actin cytoskeleton of the YSL plays critical role in epiboly (Bruce and Heisenberg, 2020). Several actomyosin driven processes during early embryogenesis, such as cytoplasmic streaming during cytoplasm and yolk separation and cortical granule exocytosis are impaired in M*dchs1b* mutants (Li-Villarreal et al., 2015), suggesting that vertebrate Dachsous cadherins are required for actomyosin cytoskeleton. Our analysis of dynamic actin organization in MZ*dchs1b* mutants during epiboly using *Tg(μ-actin:utrophin-GFP)* and Phalloidin staining showed that filamentous actin becomes enriched at the EVL and DEL margins starting at 50% epiboly in a manner comparable to WT. However, a basket-like actin meshwork of the YSL was discontinuous in the mutants, similar to the irregular microtubule network (Figure 7). Moreover, live imaging revealed that the vegetal to animal actomyosin flow in the yolk cell cortex was also impaired (Movie 1). These actomyosin flows are thought to produce friction that generates an animal-vegetal contraction that also contributes to blastoderm vegetal movement (Bruce and Heisenberg, 2020). Further investigation is necessary to determine whether the actin cytoskeletal defects are secondary to the microtubule abnormalities, or whether Dachsous regulates actomyosin cytoskeleton directly. The molecular mechanism via which Dachsous cadherins regulate actin cytoskeleton in vertebrates remains to be determined.

Our bulk transcriptomic analysis of MZ*dchs1a*;MZ*dchs1b*;MZ*dchs2* at the end of epiboly provides further insight into how these atypical cadherins regulate early development and morphogenesis. This analysis identified nearly 2,500 differentially expressed genes (DEGs) in the mutants, with most DEGs representing upregulated genes. *dchs1b* is among the downregulated genes, consistent with previous reports of nonsense mediated degradation in this mutant line (Li-Villarreal et al., 2015). Interestingly, genes associated with dorsoventral patterning are not among the DEGs. Misregulation of genes such as *goosecoid* or *chordin* was observed in some MZ*dchs1b* mutants before gastrulation and linked to abnormal transport of dorsal determinants (Li-Villarreal et al., 2015). Normalization of expression of such genes by the end of gastrulation could be due to the established regulatory capacity of the Spemann-Mangold organizer. Gene Ontology (GO) analyses point to embryonic structures including chorion, and YSL, consistent with the previously reported defects in chorion expansion upon egg activation (Li-Villarreal et al., 2015) and epiboly defects studied here. Notable predicted GO Biological Process are “actomyosin structure organization”, “actin filament organization” in line with the defects in the yolk cell actin cytoskeleton previously shown to be essential for epiboly.

### Limitations of this study

In this work we used ENU-induced frame-shift mutations that lead to premature stop codons predicted to truncate the encoded proteins in the N-terminal half. Given the large size and several predicted transcript variants of the three genes, we cannot exclude the possibility that the mutant alleles have only hypomorphic activities and that the phenotypes reported here represent the full loss of function of Dachsous cadherins.

## Declaration of interest

The authors have no financial and personal relationships with other people or organizations that could inappropriately influence their work.

## Acknowledgments

The authors would like to acknowledge members of the Solnica-Krezel laboratory for discussions, advice and assistance. We thank the Washington University Zebrafish Facility staff for their excellent zebrafish care. This research was supported in part by the grant from the National Institutes of Health R35 GM118179 to LSK and CMB training grant T32GM007067-41 to G.D.C.

## Materials and Methods

### Zebrafish lines and animal care

Zebrafish WT AB*, *dchs1a^sa15468^*, *dchs1b^fh275^*, *dchs2^stl1^*, *ttc28^stl363^*, *Tg(μ-actin:utrophin-GFP)*, *dchs1b-2xsfGFP* and *dchs1b-6xsfGFP* lines were used in this study. As described in previous studies, *dchs1b* mutant phenotypes display a female age-related decrease in penetrance and expressivity as reported for many early embryonic mutant phenotypes in zebrafish. MZ mutant embryos were generated by pairwise crossing of adult (3-12 months old) females and males homozygous for the studied mutations. In individual experiments, we used mutant and WT females of similar age. Embryos were generated by natural matings or *in vitro* fertilization and kept at 28.5°C in egg water (60μg/mL Instant Ocean in distilled water).

All zebrafish experiments and procedures were performed in accordance with the Institutional Animal Use and Care Committee of the Washington University in St. Louis School of Medicine.

### Cloning and mRNA synthesis

Human *GFP-CAMSAP2* was subcloned into the *pT7Ts* vector for mRNA synthesis and injection. All mRNAs for injections were synthesized from corresponding linearized plasmid templates using mMessage mMachine Kit (Ambion).

### Live imaging

#### Epiboly progression

Embryos were injected with 50pg of *H2B-RFP* and 200pg of *mCherry-CAAX* synthetic mRNAs at the one-cell stage to label both nuclei and cell membranes, respectively. These embryos were manually dechorionated with forceps during late cleavage stages and then mounted in 0.3% low-melting agarose on a glass bottom dish (Ted Pella, Inc.) at the 1K cell stage. Time-lapse imaging was performed beginning at the oblong stage using a spinning-disk confocal microscope (Olympus IX81, Quorum) with a 10X objective at 28.5°C. Z-stacks spanned 120μm at 1μm steps and were acquired every 10 minutes for 10 hours. Embryos requiring post-imaging genotyping were removed from the agarose the following morning.

#### EB3-GFP

Prior to the one-cell stage, embryos were injected with 50pg of synthetic mRNA encoding EB3-GFP. These embryos were mounted as described above at the 1K cell stage. Time-lapse image acquisition was performed between sphere and dome stages (4-4.5 hpf) using a Nikon spinning-disk confocal microscope with a 60X oil immersion lens at room temperature. Yolk microtubule Z-stacks spanned 4-6μm at 1-1.5μm steps. Z-stacks were acquired every two seconds.

#### Immunostaining

Immunostaining for microtubules and modified tubulin was carried out using a modified protocol from (Solnica-Krezel and Driever, 1994). Embryos at 60% epiboly stage were fixed in microtubule assembly fixative (80mM KPIPES pH 6.5, 5mM EGTA, 1mM MgCl_2_, 3.7% formaldehyde, 0.25% glutaraldehyde, and 0.2% Triton X-100 and optional 1 μM paclitaxel) for 2 hours at room temperature and then at -4°C overnight. Embryos were manually dechorionated with forceps the next day followed by overnight incubation in 100% methanol at -20°C. To quench the glutaraldehyde from the fixative, embryos were incubated in PBS containing 100mM NaBH_4_ in 24-well plastic plates for 8-10 hours at room temperature, followed by several washes in TBS and a 30-minute room temperature incubation in TBS with 2% BSA and 5% NGS. The embryos were then incubated with primary antibody (1:500 DM1A; 1:250 α-acetylated tubulin; 1:250 α-tyrosinated tubulin; 1:250 CAMSAP1 antibodies-online #ABIN2787677) in TBS containing 2% BSA and 5% NGS overnight at 4°C, followed by 5 x 10-minute washes in TBS at room temperature. The embryos were incubated in secondary antibody (1:500 Alexa Fluor 488, 568, and/or 633) in TBS containing 2% BSA and 5% NGS overnight at 4°C, followed by 5 x 20-minute washes in TBS at room temperature. DAPI staining (1:1000) was performed at room temperature for 10 min, followed by 3 x 5-minute washes in TBS. Embryos were mounted in 3% methylcellulose on a glass bottom dish (Ted Pella, Inc.) and imaged using a spinning-disk confocal microscope (Olympus IX81, Quorum) with a 10X objective collecting Z-stacks of 120μm with 1μm steps for microtubule bundling experiments, 40X objective and Z-stacks of 120μm with 1μm steps for acetylated tubulin, 60X immersion objective and Z-stacks of 60μm with 0.5μm steps for microtubule coherency experiments.

### Quantification and analysis

#### EB3-GFP quantification

To quantify microtubule polymerization track properties such as displacement, duration, and speed, EB3-GFP comets in time-lapse movies were converted to spots with 5 pixel diameters plotted at x,y positions corresponding to the centroid of each comet at each timepoint. These positions were tracked and analyzed using the TrackMate ImageJ plugin. Briefly, the difference of Gaussians detector (DoG) algorithm detected individual spots, and the Linear Assignment Problem (LAP) tracker joined these spots into tracks.

#### Tubulin modification staining quantification

To quantify the relative amount of the microtubule cytoskeleton associated with CAMSAP1 or modified tubulin, both image channels, total tubulin and the protein of interest, were thresholded using the IsoData and Moments ImageJ algorithms, respectively. All images were acquired using the same exposure, laser power, and gain settings for each channel. Together, these efforts reduced the amount of variation and human error during image acquisition and quantification. A 125x125 pixel representative square was cropped from the original image to contain only yolk microtubules located just below the YSL. Then, the channel of interest was filtered to only contain pixels that overlap with the total tubulin channel, selecting only for protein associated with microtubules. This was achieved by using the ImageJ image calculator to multiply both channels together. A pixel positive for staining has a value of 1, and any pixels negative for staining have values of 0; therefore, the only remaining pixels in the resulting image are those where both channels overlap. ImageJ was then used to calculate the total area and number of objects in the field of view.

#### MT fiber analysis

To analyze yolk microtubule fibers, a 125x125 pixel representative square was cropped from the original image to contain only yolk microtubules located just below the YSL. Orientation distributions and representative vector fields were generated using the OrientationJ plugin for ImageJ with a local window size of 2 and vector field grid size of 10 ^359^.

#### Generation of Dachsous-GFP knock-in lines

To develop Dchs1b protein reporter lines, we used a TALEN-mediated homologous recombination gene targeting technology (Shin et al., 2014). Briefly, using TALENT 2.0 (https://tale-nt.cac.cornell.edu) (Doyle et al., 2012), we designed TALENs targeting near the stop codon of dchs1b genes. The TALEN recognition sites are 5’-TCAGCCTCCATCATCAGCACC and 5’-TCTCAGCCTCATCAGTGTGCTC. The process involved in assembling RVD-containing repeats and subcloning TALE repeats into modified TALE nuclease expression vectors was performed using the REAL Assembly TALEN Kit (Addgene TALEN kit 1000000017) (Sander et al., 2011). TALEN RNAs were synthesized using SP6 mMessage mMachine Kit (Ambion) and purified by Micro Bio-SpinTM P-30 Gel columns (Bio-Rad).

To create a *dchs1b* targeting construct that is capable to induce an insertion of a long DNA fragment into the *dchs1b* stop codon site by TALEN-mediated homologous recombination, we first amplified a genomic DNA fragment (3,908 bp) containing the *dchs1b* stop codon by PCR and subcloned it into pSMART®HCKan vector (Lucigne). For the inserts, we used two copies of sfGFP (2XsfGFP) or six copies of sfGFP (6XsfGFP). With the series of cloning processes, we eventually generated three different targeting constructs (pSMART-dchs1b-2XsfGP and pSMART-dchs1b-6XsfGFP). The constructs were digested with AvrII (NEB) and the linearized targeting constructs were purified by QIAquick PCR-purification kit (Qiagen).

A pair of *dchs1b* TALEN RNAs and a targeting construct with10X injection buffer (1M KCl, 0.03% Phenol red) were mixed thoroughly and injected a final concentration of 35 ng/μL for each TALEN RNA and 10 ng/μL for the targeting construct in 1X injection buffer into the cytosol of early one-cell embryos. The injected founder pool was grown to adult stage and F1 progeny were screened by PCR using sfGFP-specific primers. All insertion-positive embryos were confirmed by additional PCR-based genotyping and Sanger sequencing to further confirm that the insert was integrated into the target locus without error.

### Statistical analysis

All statistical analyses were completed using GraphPad Prism 7. Statistical significance was calculated using a two-tailed unpaired Student’s *t*-test to compare two populations and Kolmogorov-Smirnov tests were used to compare population distributions between two groups.

### RNA-seq

Embryos were maternal zygotic for dchs1b, dchs1a, and dchs2 (MZdcsh1a; MZdcsh1Bfh2 MZdchs2ALLELES). “WT” RNA came from an AB* stock. RNA was extracted from embryos at 10 hpf using Trizol (Invitrogen). Pooled RNA was treated with DNAse, and cleaned up on Qiagen RNEasy mini kit columns. Approximately 1 ug of RNA was submitted to the Genome Technology Access Center at the McDonnell Genome Institute at Washington University School of Medicine. Total RNA integrity was determined using Agilent 4200 Tapestation. Library preparation was performed with 1μg of total RNA. Ribosomal RNA was removed by a hybridization method using Ribo-ZERO Gold kits (Illumina-EpiCentre). mRNA was then fragmented in reverse transcriptase buffer and heating to 94 degrees for 8 minutes. mRNA was reverse transcribed to yield cDNA using SuperScript III RT enzyme (Life Technologies, per manufacturer’s instructions) and random hexamers. A second strand reaction was performed to yield ds-cDNA. cDNA was blunt ended, had an A base added to the 3’ ends, and then had Illumina sequencing adapters ligated to the ends. Ligated fragments were then amplified for 15 cycles using primers incorporating unique dual index tags. Fragments were sequenced on an Illumina NovaSeq-6000 using paired end reads extending 150 bases. Samples were prepared according to library kit manufacturer’s protocol, indexed, pooled, and sequenced on an Illumina NovaSeq 6000.

### RNA-seq data analyses

Reads were processed using an in-house pipeline and open-source R packages. Briefly, raw reads were first trimmed using cutadapt to remove low quality bases and reads. Trimmed reads were then aligned to the zebrafish genome GRCz11 with ENSEMBL annotation using STAR (v2.7.9a) with default parameters. Transcript quantification was performed using featureCounts from the subread package (v2.0.1). further quality control assessments were made using RSeQC and RSEM, and batch correction was performed using edgeR, EDASeq, and RUVSeq. Principal component analysis (PCA) and differential expression analysis for WT, Mild, Severe MZ*dchs* triple mutant groups were determined using DESeq2 in negative binomial mode using batch-corrected transcripts from featureCounts (> 2-fold expression change, > 1 count per million (CPM), Benjamini corrected P < 0.05). Pairwise comparisons were made between groups to determine differentially expressed genes (DEGs). Gene ontology (GO) and KEGG analyses were performed using FishEnrichr. The gene expression was plotted using ggplot2 and pheatmap. Comparison of lists of DEGs and Venn Diagrams were prepared at molbiotools.com (https://molbiotools.com/listcompare.php).

### Method details

#### EB3-GFP dynamics imaging

Embryos were injected with 50pg of RNA encoding EB3-GFP prior to the one-cell stage. Injected embryos were mounted as described above at the 1,000-cell stage and image acquisition was performed between sphere and dome stages (4 – 4.5 hpf). Time-lapse imaging was performed using a Nikon spinning-disk confocal microscope with a 60X oil immersion lens at room temperature. Z-stack was set up for a total of 4-6 μm at 1-1.5 μm interval. Images were acquired every 2 seconds.

#### EB3-GFP tracks analysis

To quantify the speed, duration, and displacement parameters of individual EB3-GFP tracks, comets in time-lapses were converted to spots with a diameter of 5 pixels plotted at x,y positions corresponding to the centroid of each comet at each timepoint. These positions were tracked and analyzed using the TrackMate ImageJ plugin.^350^ The difference of Gaussians detector (DoG) algorithm was used to detect individual spots, and the linear motion Linear Assignment Problem (LAP) tracker was used to join these spots into tracks. To quantify trajectory angles of individual EB3-GFP tracks, angles were calculated between each comet’s initial and final locations.

#### Statistical analyses

All statistical analyses were performed with PAST and Graphpad Prism 7 software. For midzone microtubule bundling experiments, paired Student’s two-tailed t-test was used to determine the difference between control and mutants. See figure legends for additional information about number of embryos used in each experiment.

---

**Supplement Figure 1.**
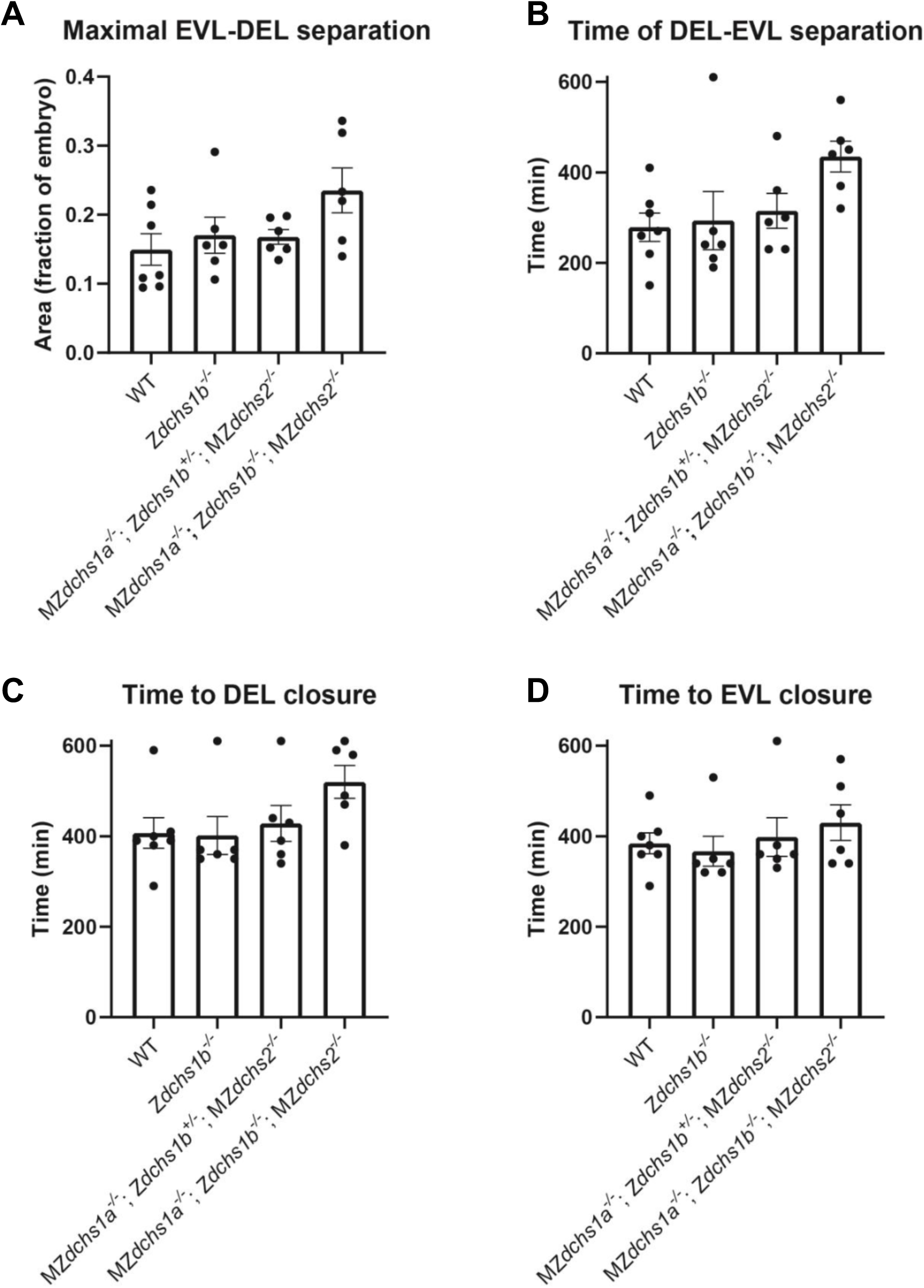
Additional epiboly progression quantification in wild-type and *dchs* triple mutant embryos, relevant to Figure 1. A. Quantification of maximal YSL-DEL separation during epiboly. Kruskal-Wallis test, multiple comparisons, ns. B. Quantification of the elapsed time of YSL-DEL separation during epiboly. Kruskal-Wallis test, multiple comparisons, ns. C. Quantification of elapsed time for DEL closure. Kruskal-Wallis test, multiple comparisons, ns. D. Quantification of elapsed time for EVL closure. Kruskal-Wallis test, multiple comparisons, ns.

**Supplement Figure 2.**
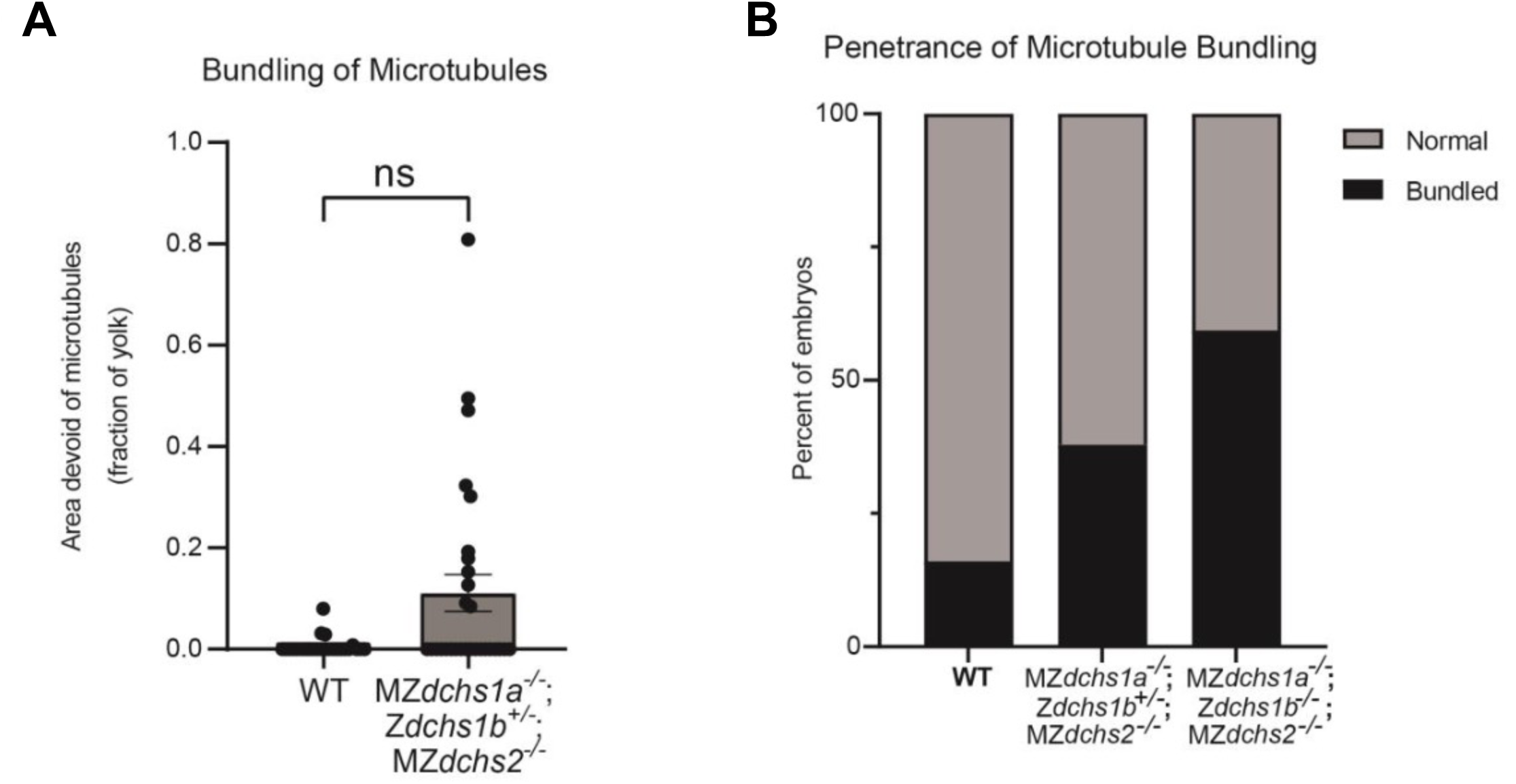
Expressivity and penetrance of microtubule bundling in compound *dchs* triple mutants, relevant to Figure 3. A. Quantification of yolk microtubule bundling through measurement of bare yolk area lacking microtubules at 50% epiboly. ns (p=0.0887). WT N=25 embryos, MZ*dchs1a^-/-^*; Z*dchs1b^+/-^*; MZ*dchs2^-/-^* N=29 embryos. B. Quantification of number of embryos with yolk microtubule bundling. WT N=25 embryos, MZ*dchs1a^-/-^*; Z*dchs1b^+/-^*; MZ*dchs2^-/-^*N=29 embryos, MZ*dchs1a^-/-^*; Z*dchs1b^-/-^*; MZ*dchs2^-/-^* N=32 embryos.

**Supplement Figure 3.**
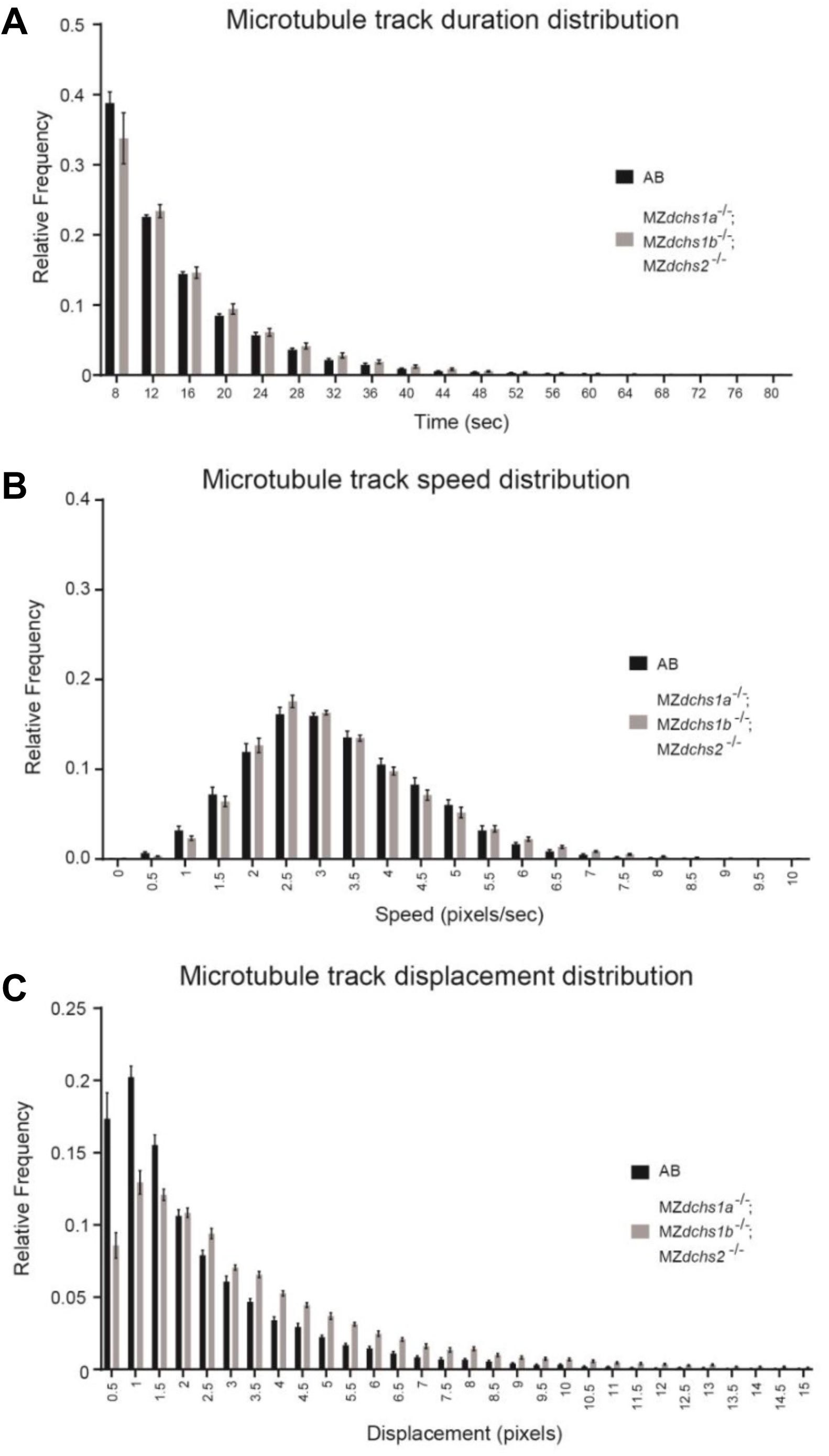
Microtubule polymerization dynamics in MZ*dchs* triple mutants. A. Frequency distribution of EB3 track duration. Error bars indicate SEM. WT N=9 embryos. MZ*dchs1a^-/-^*; MZ*dchs1b^-/-^*; MZ*dchs2^-/-^* N=14 embryos. Wilcoxon matched-pairs rank test, **p<0.01 B. Frequency distribution of EB3 track average speed. Error bars indicate SEM. WT N=9 embryos. MZ*dchs1a^-/-^*; MZ*dchs1b^-/-^*; MZ*dchs2^-/-^* N=14 embryos. Wilcoxon matched-pairs rank test, ns C. Frequency distribution of EB3 track displacement. Error bars indicate SEM. WT N=9 embryos. MZ*dchs1a^-/-^*; MZ*dchs1b^-/-^*; MZ*dchs2^-/-^* N=14 embryos. Wilcoxon matched-pairs rank test, **p<0.01

**Supplement Figure 4.**
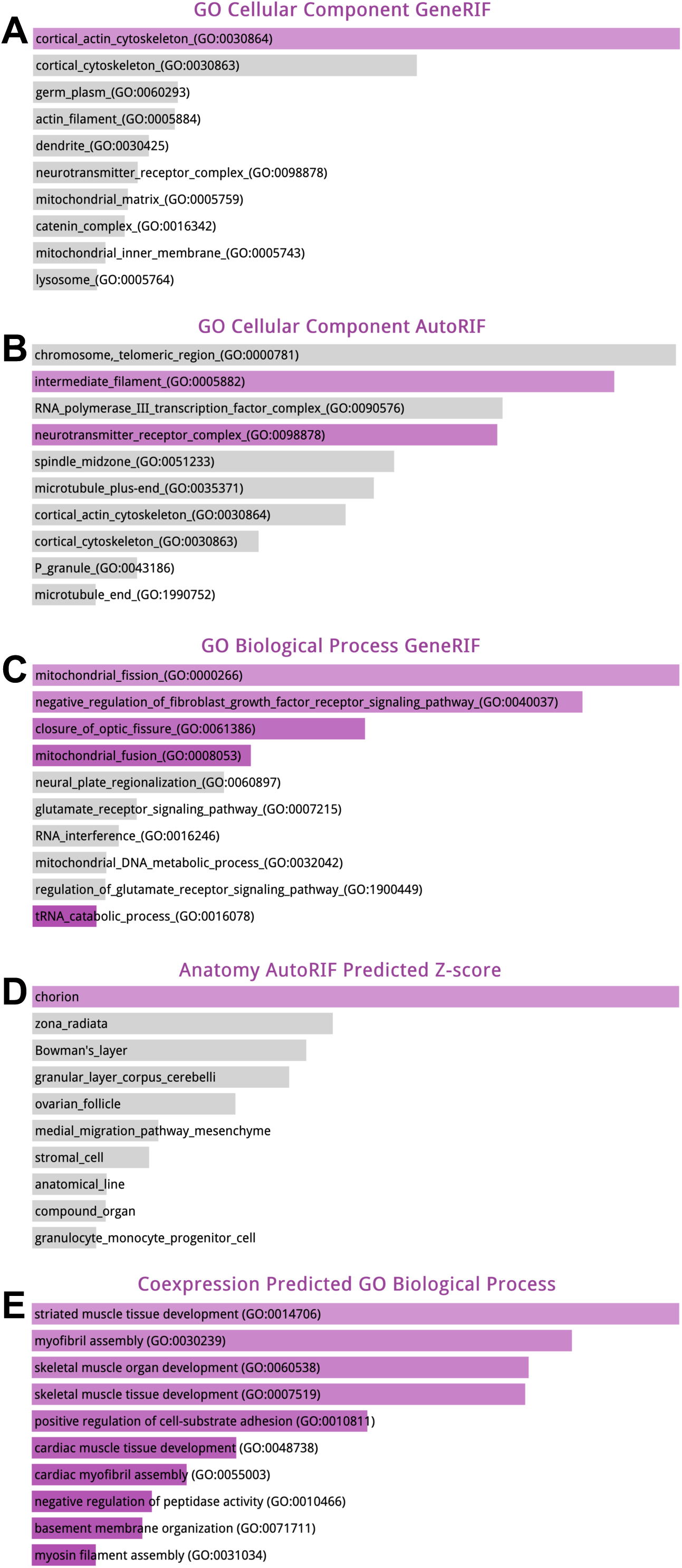
FishEnricher pathway analysis of 781 genes downregulated in MZ*dchs1a,b,2* triple mutants (A-D). FishEnricher pathway analysis of 781 genes upregulated in MZ*dchs1a,b,2* triple mutants (E)

**Supplemental Movie 1.**
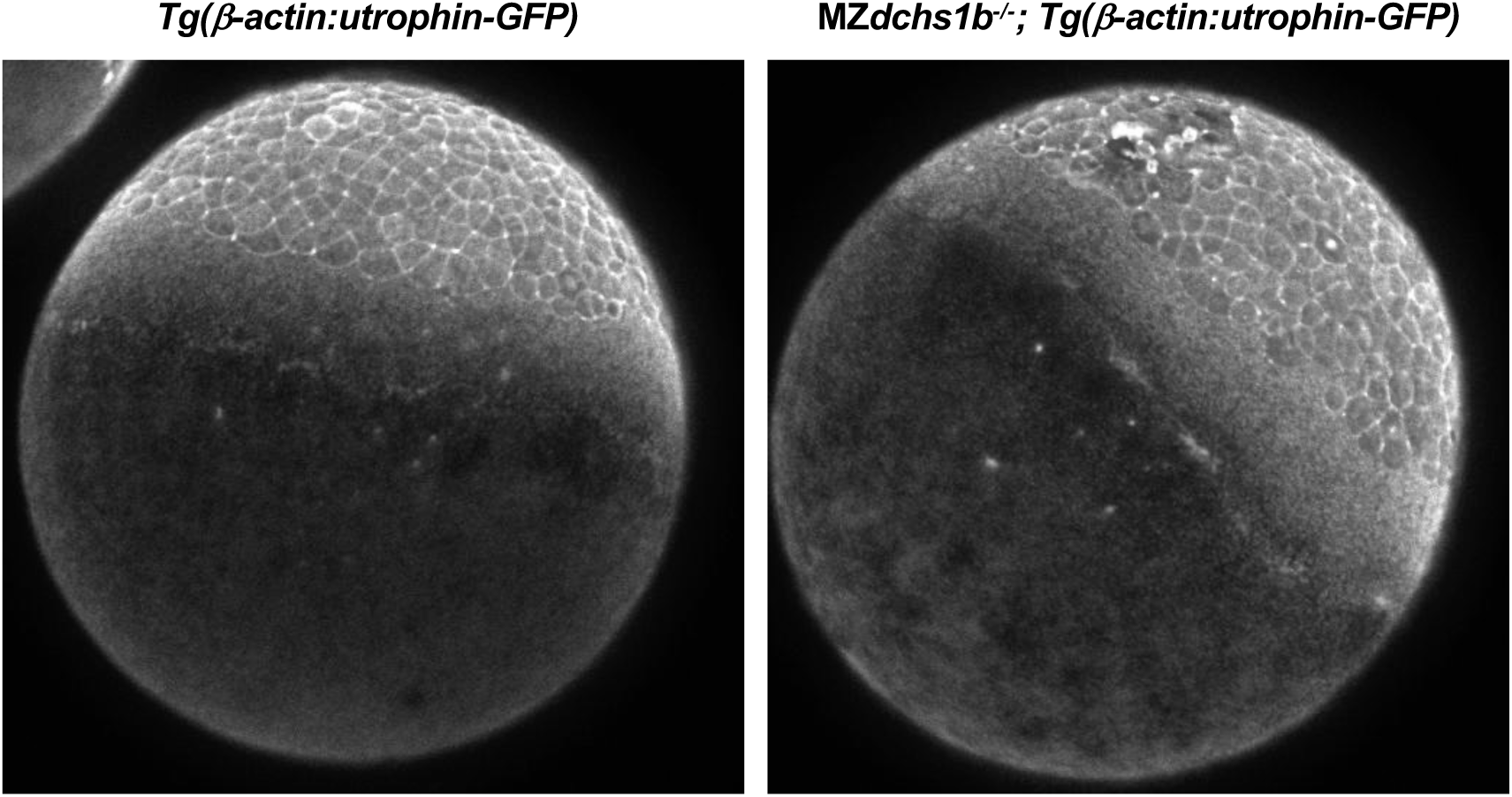
Supplemental Movie in support of Figure 7. *Tg(m-actin:utrophin-GFP)* in control WT and MZ*dchs1b* mutant embryo imaged in lateral view. In WT the animal pole is at the top and in the mutant, it is tilted to the right. Time-lapse imaging was conducted at 23°C from ∼sphere stage, the onset of epiboly, and continued until epiboly was completed in the WT embryo.

## Notes

### Competing Interest Statement

The authors have declared no competing interest.

