## Supplementary figures and images for "Regulation of the Yolk Microtubule and Actin Cytoskeleton by Dachsous Cadherins during Zebrafish Epiboly"

### Supplemental Movie1

#### Slide 1
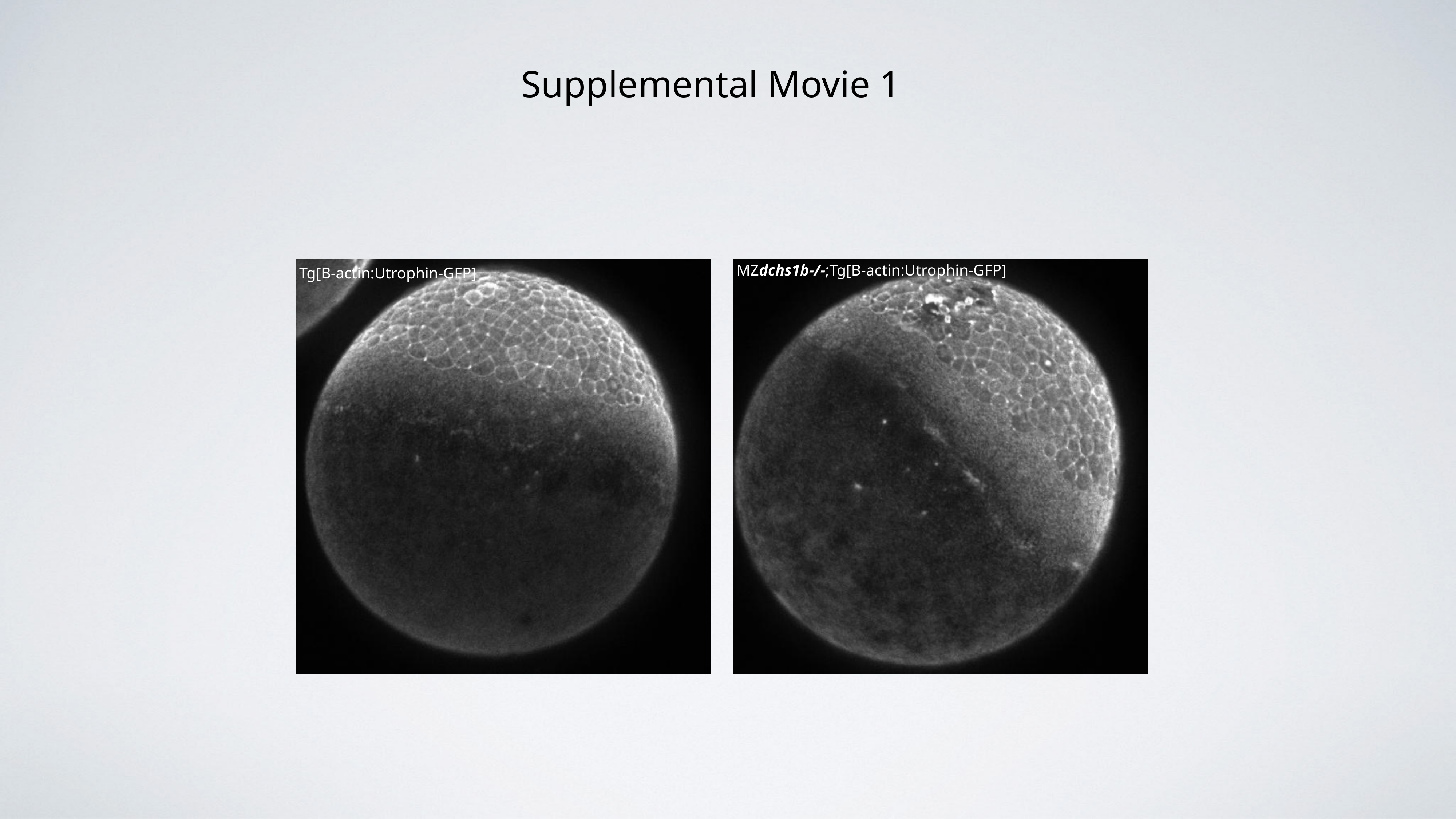

Supplemental Movie 1
MZdchs1b-/-;Tg[B-actin:Utrophin-GFP]
Tg[B-actin:Utrophin-GFP]
